# ORP5 controls the partitioning of phosphatidic acid between triacylglycerol and cardiolipin synthesis at mitochondria–ER–lipid droplet contact sites

**DOI:** 10.1101/2025.11.06.685814

**Authors:** Vera F. Monteiro-Cardoso, Valentin Guyard, Jennica Trager, Cécile Sauvanet, Helin Elhan, Mehdi Zouiouich, Naima El Khallouki, Yoshiki Yamaryo-Botté, David Tareste, Cyrille Y. Botté, Abdou Rachid Thiam, Francesca Giordano

## Abstract

Phosphatidic acid (PA) is a central metabolic intermediate that can fuel triacylglycerol (TAG) synthesis in lipid droplets (LDs) or cardiolipin production in mitochondria, but how cells partition PA between these competing fates has remained a fundamental unresolved question in lipid cell biology. We identify the lipid transfer protein ORP5 as a key regulator of PA partitioning at Mitochondria–Associated endoplasmic reticulum Membranes (MAM) that contact lipid droplets (LD), referred to as MAM–LD junctions. Cell imaging analysis shows that ORP5 stabilizes PA levels at MAM to promote TAG synthesis. On the other hand, loss of ORP5 causes PA accumulation on mitochondrial membranes, leading to excess cardiolipin synthesis and mitochondrial hyperfusion, while impairing triacylglycerol (TAG) synthesis and LD formation. Finally, reconstitution assays using liposomes or giant organelles further demonstrate that ORP5 can transfer PA from mitochondria to the ER via its ORD domain. Together, these findings reveal that ORP5 functions as a PA lipid transfer protein at tripartite MAM–LD contacts, where it balances LD formation with mitochondrial lipid metabolism, protecting mitochondria from cardiolipin overload.

## INTRODUCTION

Eukaryotic cells maintain their energy balance by finely tuning lipid synthesis, storage and metabolism. Lipid droplets (LDs) act as the primary reservoirs of neutral lipids, buffering energy fluctuations and protecting against lipotoxic stress. At the same time, mitochondria convert lipids into metabolic energy and use specific lipid precursors for the synthesis of essential phospholipids (Decker and Funai 2024; Thiam, Farese, and Walther 2013; Olzmann and Carvalho 2019). These organelles are therefore central hubs of lipid and energy homeostasis: LDs sequester fatty acids as triacylglycerol (TAG) and sterol esters within a phospholipid monolayer, whereas mitochondria oxidize fatty acids within their cristae membranes to fuel ATP production (Enkler and Spang 2024). The physical and metabolic coupling between LDs and mitochondria is thus vital for cellular and systemic metabolic balance, particularly in energy-demanding tissues such as liver, muscle, and adipose tissue (Klemm and Carvalho 2024). Dysregulation of this crosstalk contributes to obesity, hepatic steatosis, and mitochondrial dysfunction (Herker et al. 2021; Klein et al. 2022; Suomalainen and Nunnari 2024).

The endoplasmic reticulum (ER), the main site of lipid synthesis, supplies both mitochondria and LDs with key lipid precursors and intermediates for lipid synthesis. Mitochondria depend on imported phospholipids from the ER, such as phosphatidylserine (PS) and phosphatidic acid (PA), to generate phosphatidylethanolamine (PE) and cardiolipin, which are essential for mitochondrial cristae structure and respiratory function (Vance 2015; Horvath and Daum 2013; Giordano 2018). LDs originate from the ER, where neutral lipids accumulate within the bilayer at sites marked by seipin, a scaffolding protein essential for LD nucleation (Salo et al. 2019; Thiam and Ikonen 2021; Salo 2023). Furthermore, the ER provides LDs with many of their monolayer phospholipids and with precursors for neutral lipid synthesis, mainly TAG and sterol esters. Yet, how lipids are delivered to LDs during their formation and growth remains a fundamental open question.

Because neither LDs nor mitochondria integrate into the classical vesicular transport pathways, non-vesicular lipid transfer between them and with the ER relies on membrane contact sites (MCS), zones of close apposition between organelles without membrane fusion (Scorrano et al. 2019), and on lipid transfer proteins (LTPs) localized at these sites (Prinz, Toulmay, and Balla 2020; Voeltz et al. 2024). Major contacts include ER–mitochondria, ER– LD, and mitochondria–LD (also termed peridroplet mitochondria in adipocytes) (Herrera-Cruz and Simmen 2017; Benador et al. 2018; Hugenroth and Bohnert 2020). Despite recent progress in identifying novel components and LTPs at these contacts (Wong, Gatta, and Levine 2019), the molecular mechanisms that govern lipid flux across them remain only partially defined, and the repertoire of LTPs operating at these interfaces is still largely unknown, emphasizing the need to further delineate the principles of lipid partitioning and organelle communication.

We recently showed that mitochondria-associated ER membranes (MAM) are key sites for LD biogenesis and growth, identifying the existence of tripartite mitochondria-ER-LD junctions (MAM-LD contacts), that act as metabolic hubs for lipid transfer and storage (Monteiro-Cardoso and Giordano 2024; Guyard et al. 2022). Together with the recent discoveries of three-way organelle junctions (Joshi et al. 2018; Elbaz-Alon et al. 2020; Guillen-Samander et al. 2021; Guyard and Giordano 2025), these findings underscore that lipid metabolism occurs within an intricate network of inter-organelle communication. Yet, many questions remain unresolved: which lipid species are exchanged at MAM–LD contacts, how fluxes are partitioned between storage in LDs and utilization in mitochondria, which LTPs orchestrate these processes.

Among candidate lipid intermediates, PA occupies a central position(Athenstaedt and Daum 1999). On one hand, it serves as the entry point into TAG synthesis and LD expansion. On the other hand, PA is the precursor for mitochondrial cardiolipin biosynthesis (Zhang and Reue 2017). Beyond its metabolic role, PA also modulates membrane curvature and can serve as a signaling lipid that recruits protein complexes to MCS (Kim et al. 2015; Chang and Liou 2015; Yan et al. 2018; Thakur et al. 2019; Zhou et al. 2024). Despite these critical functions, it remains unclear how PA flux is spatially regulated at MAM–LD junctions to coordinate LD expansion with mitochondrial lipid demands.

Members of the oxysterol-binding protein (OSBP)-related protein (ORP) family are emerging as central regulators of lipid exchange at MCS (Olkkonen 2015; Nakatsu and Kawasaki 2021). ORPs share a conserved OSBP-related domain (ORD) capable of lipid binding and transfer (de Saint-Jean et al. 2011; Mesmin et al. 2013; Pietrangelo and Ridgway 2018). Most also carry a pleckstrin homology (PH) domain and an FFAT motif, enabling dual targeting to the ER (via VAP proteins) and partner membranes (via phosphoinositides). ORP5 and ORP8 are unique in that they lack an FFAT motif but contain a hydrophobic stretch anchoring them to the ER, and a PH domain that binds plasma membrane phosphoinositides, especially PI(4)P and PI(4,5)P_2_ (Chung et al. 2015; Ghai et al. 2017).

Initially characterized at ER–plasma membrane contacts as mediators of PS/PI(4)P exchange (Chung et al. 2015), ORP5 and ORP8 were later found to localize predominantly at ER–mitochondria contacts, where they mediate PS transfer for mitochondrial PE synthesis and regulate LD biogenesis at MAM–LD sites (Galmes et al. 2016; Guyard et al. 2022; Monteiro-Cardoso et al. 2022). At these sites, ORP5/8 also interact with seipin (Guyard et al. 2022). Seipin itself localizes to ER–mitochondria contacts, where it regulates calcium uptake and bioenergetics, further linking lipid storage to mitochondrial function (Combot et al. 2022). While ORP5/8 are established PS transfer proteins, whether they also regulate the flux of other phospholipids has remained unclear. Notably, MAM subdomains enriched in ORP5 contain high levels of PA, and seipin can bind anionic lipids including PA (Yan et al. 2018). Recent work has further implicated seipin in PA handling (House et al. 2025). Lastly, we found that the CC domain of ORP5 binds to PA *in vitro* (Guyard et al. 2022). While all this data suggests potential localized PA metabolism at the LD-mitochondria interface, it remains an open question whether ORP5 regulates PA distribution between these organelles, linking TAG synthesis to mitochondrial lipid homeostasis.

In this study, we identify ORP5 as a key regulator of PA metabolism at MAM–LD contacts. Using a combination of imaging, biochemical, lipidomic, *in vitro*, and *ex vivo* assays, we show that ORP5 stabilizes PA at ER-mitochondria subdomains and preferentially favors its transfer from mitochondria toward the associated ER for TAG synthesis. Loss of ORP5 alters mitochondrial PA levels, leading to increased cardiolipin synthesis and mitochondrial hyperfusion, while ORP5 overexpression promotes TAG synthesis and LD growth. Our findings reveal a previously unrecognized role for ORP5 in balancing lipid storage with mitochondrial lipid biosynthesis, highlighting PA transport at the MAM–LD junctions as a key mechanism of inter-organelle metabolic coordination.

## RESULTS

### ORP5 is required for efficient PA conversion in TAG and storage in lipid droplets

We have previously shown that the ER-anchored lipid transfer protein ORP5 regulates LD biogenesis at MAM and that this function depends on its ORD domain, implicating ORP5 lipid transfer activity in LD biogenesis (Guyard et al. 2022). However, the specific lipids involved remain unknown. ORP5 is established as a PS lipid transfer protein at ER-mediated contact sites with other organelles, including LDs (Du et al. 2020), raising the possibility that the regulation of PS distribution by ORP5 could be involved in LD biogenesis. Unexpectedly, expression of the ORP5(L489D) mutant, which is defective in PS transport (Chung et al. 2015), rescued the LD biogenesis defects caused by ORP5 depletion (Supp. Fig. 1). This indicates that PS transfer is not required for ORP5-mediated LD formation.

These findings led us to test whether ORP5 regulates other lipids critical for LD growth. Given the central role of PA as the immediate precursor for TAG synthesis (Fig. 1A), we examined PA distribution in HeLa cells using a fluorescent PA analog (TopFluor-TMR-PA) (Zhang and Reue 2017). Fluorescent lipid derivatives with modified fatty acyl chains are established probes for monitoring lipid trafficking and metabolism (Kay et al. 2012; Zewe et al. 2020; Niu et al. 2024). We first monitored the incorporation of exogenous TopFluor-TMR-PA in live wild-type (*WT)* and ORP5 knockout (*KO*) HeLa cells. Interestingly, we observed that over time, TopFluor-TMR-PA accumulates in the LDs core. Notably, the accumulation of PA inside the LDs was stronger in *WT* cells than in *ORP5KO* cells (Supp. movies 1A-D and Supp. Fig. 2). To confirm this observation, we assessed the intensity of TopFluor-TMR-PA-derived signal co-localizing with LDs in fixed cells, which have been treated with TopFluor-TMR-PA for 30 minutes. To minimize possible variability in PA uptake between *WT* and ORP5-depleted cells, the intensity of TopFluor-TMR-PA in the LD was normalized by its mean intensity in the whole cell. Consistent with the observation in live cells, the loss of ORP5 in *ORP5KO* cells significantly reduced TopFluor-TMR-PA-derived signal inside LDs as compared to *WT* cells (Fig. 1B,C). Additionally, *ORP5KO* cells loaded with TopFluor-TMR-PA displayed a significant reduction in LD number and volume (Fig. 1E-F). Importantly, regression analysis revealed that TopFluor-TMR-PA incorporation into LD occurred independently of LD size under both conditions (Fig. 1F).

**Figure 1:**
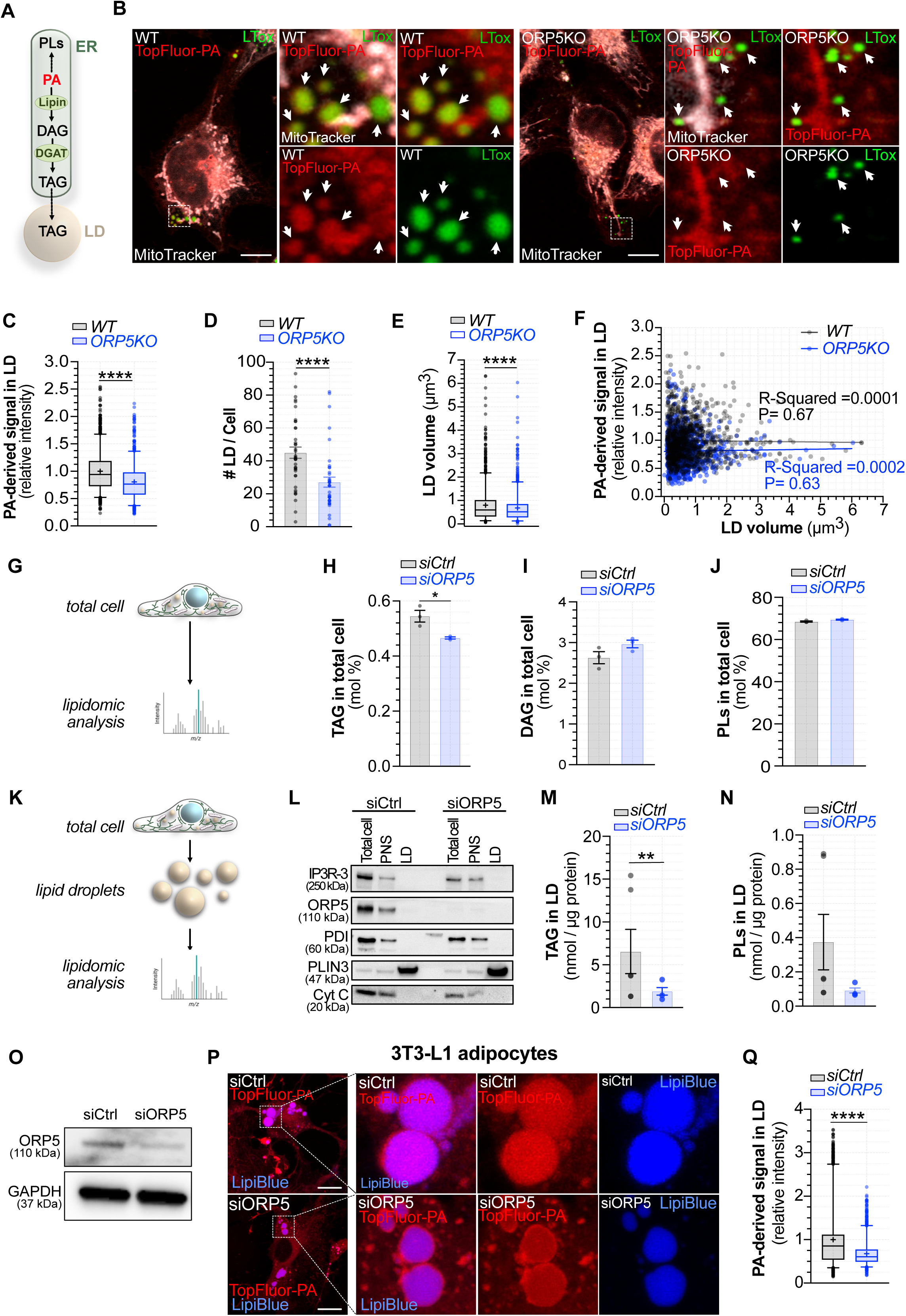
ORP5 facilitates the metabolism of phosphatidic acid into triacylglycerols and its storage in lipid droplets. **(A)** Schematic representation of phosphatidic acid (PA) metabolism. PA is a metabolic intermediate for phospholipids (PLs) and triacylglycerols (TAG). The synthesis of TAG in the endoplasmic reticulum (ER) is mediated by the enzymatic processing of PA by Lipin, which generates diacylglycerol (DAG), followed by DGAT. TAG fuels lipid droplets biosynthesis and growth. **(B)** Representative confocal images, including full cell and zoomed regions, of wild-type (*WT)* and ORP5 knockout (*ORP5KO)* HeLa cells treated with 2 µM TopFluor-TMR-PA (red) for 30min and stained with MitoTracker deep red (gray) and LipidTox (purple) to mark mitochondria and lipid droplets, respectively. Scale bar, 10 µm. **(C)** Relative quantification TopFluor-TMR-PA-derived fluorescence intensity co-localizing with lipid droplets in *WT* and *ORP5KO* cells. The absence of ORP5 reduces the accumulation of TopFluor-TMR-PA in lipid droplets in HeLa cells. Data are represented as boxplots showing the interquartile range, the median and the mean (+) of *n* = 889 - 1295 lipid droplets in three independent experiments. Statistical significance between data sets were assessed using using Mann-Whitney U test, with ****p < 0.0001. **(D)** Number of lipid droplets per cell in *WT* and *ORP5KO* HeLa cells treated with TopFluor-TMR-PA. ORP5KO displays a reduced number of lipid droplets per cell. Data are represented as mean ± SEM of *n* = 39 - 40 cells. Statistical significance between *WT* and *ORP5KO* were assessed using Mann-Whitney U test, with ****p < 0.0001. **(E)** Lipid droplets volume in WT and ORP5 treated with TopFluor-TMR-PA. *ORP5KO* cells display smaller lipid droplets than *WT* cells. Data are represented as boxplots showing the interquartile range, the median and the mean (+) of *n* = 889 - 1295 lipid droplets from three independent experiments. Statistical significance between data sets were assessed using using Mann-Whitney U test, ****p < 0.0001. **(F)** Scatter plot of TopFluor-TMR-PA derived signal intensity and lipid droplets volume in *WT* and *ORP5KO* cells. Data points, 889 - 1295 lipid droplets, were fitted with simple linear regression with P value and R-squared indicated in the plots. Top-Fluor-TMR-PA accumulation in the lipid droplets is independent of the lipid droplets size**. (G)** Simplified illustration of conceptual framework of total cell lipidomic analysis. **(H)** Triacylglycerol (TAG) content in lipid extract isolated from HeLa cells 48 h after transfection with control siRNA (*siCtrl*) or with a *ORP5*-specific siRNA (*siORP5*). Depletion of ORP5 results in decreased levels of TAG in the total cell lipid extracts. **(I)** Diacylglycerol (DAG) content in lipid extract isolated from untreated control HeLa cells (*siCtrl*) and treated with RNAi targeting ORP5 (*siORP5*). **(J)** Phospholipids (PLs) content in lipid extract isolated from untreated control HeLa cells (*siCtrl*) and treated with RNAi targeting ORP5 (*siORP5*). **(K)** Simplified illustration of conceptual framework of lipid droplets lipidomic analysis. **(L)** Immunoblotting showing reduced expression of ORP5 via RNAi treatment and lipid droplets purity. PDI, ORP5, IP3R-3 are makers for ER and MAM markers and r cytochrome C (Cyt C) is a mitochondrial marker. **(M)** Triacylglycerol (TAG) content in lipid extract obtained from lipid droplets isolated from *siCtrl* and *siORP5* HeLa cells treated with 250 µM oleic acid overnight. Depletion of ORP5 results in decreased levels of TAG in lipid droplets. **(N)** Phospholipids (PLs) content in lipid extract obtained from lipid droplets isolated from control HeLa cells (*siCtrl*) and treated with RNAi targeting ORP5 (*siORP5*) treated with 250 µM oleic acid overnight. **(O)** Immunoblotting showing the efficacy of RNAi targeting ORP5 in 3T3-TL1 ten days-post differentiation. **(P)** Illustrative confocal images, including full cell and zoomed regions, of 3T3-L1 adipocytes control and lacking ORP5, 10 day-post-differentiation, treated with 2 µM TopFluor-TMR-PA (red) for 30min and with LipiBlue (blue) to stain lipid droplets. Scale bar, 10 µm. **(Q)** Relative quantification TopFluor-TMR-PA-derived fluorescence intensity co-localizing with lipid droplets in control and ORP5-depleted 3T3-TL1 ten days-post differentiation. The absence of ORP5 reduces the accumulation of TopFluor-TMR-PA in lipid droplets in 3T3-TL1 adipocytes. Data are represented as boxplots showing the interquartile range, the median and the mean (+) of *n* = 1407 - 1166 lipid droplets from three independent experiments. Statistical significance between data sets was assessed using Mann-Whitney U test, with ****p < 0.0001. Data sets in H,J, M, N and M are represented as mean ± SEM of at least three independent experiments. *siCtrl* and *siORP5* groups were compared with Student’s t test, *p < 0.05, **p < 0.01.

To exclude the possibility that this phenotype reflects long-term adaptation to ORP5 loss, we performed similar experiments after RNAi-mediated knockdown (KD) of ORP5. Efficient depletion of ORP5 was confirmed by immunoblotting (Suppl. Fig. 3A). Acute KD phenocopied the KO, reducing PA incorporation into LDs (Supp. Fig. 3B,C), supporting that this effect is a direct consequence of ORP5 loss, and not merely an adaptive response to chronic KO.

As the core of LDs is primarily composed of TAG and cholesterol esters and can harbor DAG, we next investigated whether ORP5 could also contribute to cholesterol storage. *WT* and *ORP5KO* cells were treated with TopFluor-cholesterol. As expected, TopFluor-cholesterol localized to LDs (Supp. Fig. 3D), but its incorporation was unaffected by ORP5 depletion (Supp. Fig. 3E). These findings suggest that ORP5 selectively regulates PA storage, without directly impacting cholesterol ester accumulation.

To test whether the observed effect on TopFluor-TMR-PA dynamics and incorporation into LDs was dependent on lipid storage capacity, we preloaded *WT* and *ORP5KO* cells with oleic acid to promote LD formation. Even under these conditions, ORP5 loss significantly reduced TopFluor-TMR-PA incorporation into LDs, regardless of LD size, and decreased LD number (Supp. Fig. 3F-J), further supporting a specific role of ORP5 in channeling PA toward TAG synthesis and lipid storage, independent of LD abundance and size. Importantly, when living cells were subjected to hypotonic swelling to improve visualization of organelle interfaces, *ORP5KO* cells, preloaded with oleic acid to promote LD formation, again showed a pronounced decrease in PA signal within LDs relative to the ER (Supp. Fig. 3L-M), validating the defect under conditions where ER–LD contacts are more easily resolved.

To directly evaluate whether ORP5 influences cellular TAG metabolism, we performed mass-spectrometry-based lipidomics on total cell lipid extracts from *WT*, *ORP5KO*, and *ORP5KD* cells. ORP5-deficient (KO and KD) HeLa cells displayed reduced TAG levels (Fig. 1G, H; Suppl. Fig. 3K), while levels of diacylglycerol (DAG), the immediate TAG precursor, remained unchanged (Fig. 1I). Because PA also serves as a central intermediate for phospholipid biosynthesis, we next quantified total phospholipid levels in ORP5-deficient cells. We found that phospholipid abundance was comparable to that of control cells (Fig. 1J; Suppl. Fig. 3K). These data indicate that ORP5 specifically promotes the conversion of PA into TAG for lipid storage, without broadly perturbing phospholipid metabolism.

We next confirmed that the decrease in TAG occurred specifically in LDs. Lipid droplets were isolated from oleic acid–treated *ORP5KD* cells, and the levels of TAG and PLs were analyzed by mass spectrometry (Fig. 1K). Purity of LD preparations was validated by Western blot. LD fraction was enriched in the LD marker perilipin 3 (PLIN3) and free of specific ER and MAM (PDI, ORP5, IP3R-3) markers, as well as the mitochondria marker cytochrome C (Cyt C) (Fig. 1L). Isolated LDs from *ORP5KD* cells contained significantly less TAG (Fig. 1M), whereas phospholipid content showed a slight non-significant decrease (Fig. 1N). These results corroborate our imaging data, demonstrating that ORP5 deficiency reduces both the number and size of LDs due to impaired PA metabolism and subsequent TAG storage.

Considering the crucial role of lipid metabolism in adipocytes, we performed additional experiments in 3T3-L1 pre-adipocytes, which were differentiated into mature adipocytes (Suppl. Fig. 4A,B), to evaluate the physiological significance of our findings in HeLa cells. We also reduced ORP5 levels using siRNAs. To monitor adipocyte differentiation, we measured the mRNA expression of PPARγ (Suppl. Fig. 4C), which consistently increased through the differentiation (Suppl. Fig. 4C). Additionally, we assessed ORP5 and ORP8 protein levels (Suppl. Fig. 4D,E). Interestingly, ORP8 expression levels were not strongly impacted during adipocyte differentiation, while ORP5 expression was highly variable. During differentiation, ORP5 protein expression was initially high, which would be consistent with a role in LD biogenesis at MAMs and in PA channeling into TAG. In line with this, we found that ORP5, and to a lesser extent ORP8, at endogenous levels localize simultaneously in close proximity with mitochondria and LDs (Suppl. Fig. 4F). The presence of ORP5 at MAM-LD associations in 3T3-L1 was further confirmed in conditions where we overexpressed EGFP-ORP5 (Suppl. Fig. 4G). Similarly to what we have previously reported in HeLa cells, in 3T3-L1 adipocytes at 7 days of differentiation, EGFP-ORP5A positive ER subdomains wrap the LDs in regions where it is often associated with mitochondria.

After differentiation, ORP5 KD efficiency was confirmed by WB (Fig. 1O). Control and ORP5-depleted adipocytes were then incubated with TopFluor-TMR-PA and stained with LipiBlue to visualize LDs. Consistent with our findings in HeLa cells, depletion of ORP5 markedly reduced TopFluor-TMR-PA-derived signal within LDs (Fig. 1P,Q), supporting a conserved role for ORP5 in promoting PA storage as TAG. Together, these results suggest that ORP5 contributes to LD maturation and lipid storage in differentiated adipocytes, acting in a complementary manner to seipin.

### ORP5 regulates PA storage in lipid droplets independently of ORP8 and seipin

To further investigate the role of ORP5 in PA metabolism, we monitored the incorporation of exogenous PA into LDs using TopFluor-TMR-PA under overexpression of different ORP5 constructs in fixed cells. We expressed EGFP-tagged ORP5A, ORP5B, ORP5ΔCC or the ER-resident Sec22b, together with markers for mitochondria (Mito-BFP) and LDs (Lipid Tox). ORP5A corresponds to the full-length protein that localizes to both ER-PM and ER-mitochondria contacts (Chung et al. 2015; Galmes et al. 2016). ORP5B lacks the PH domain required for PI(4)P/ PI(4,5)P_2_ binding at PM (Du et al. 2011; Ghai et al. 2017; Guyard et al. 2022), while ORP5ΔCC is deficient in the coiled-coil (CC) domain required for ORP5-ORP8 interaction (Guyard et al. 2022) (Fig. 2A). By comparing these variants, we sought to determine which ORP5 domains contribute to PA stabilization and partitioning at MAM-LD contact sites.

**Figure 2:**
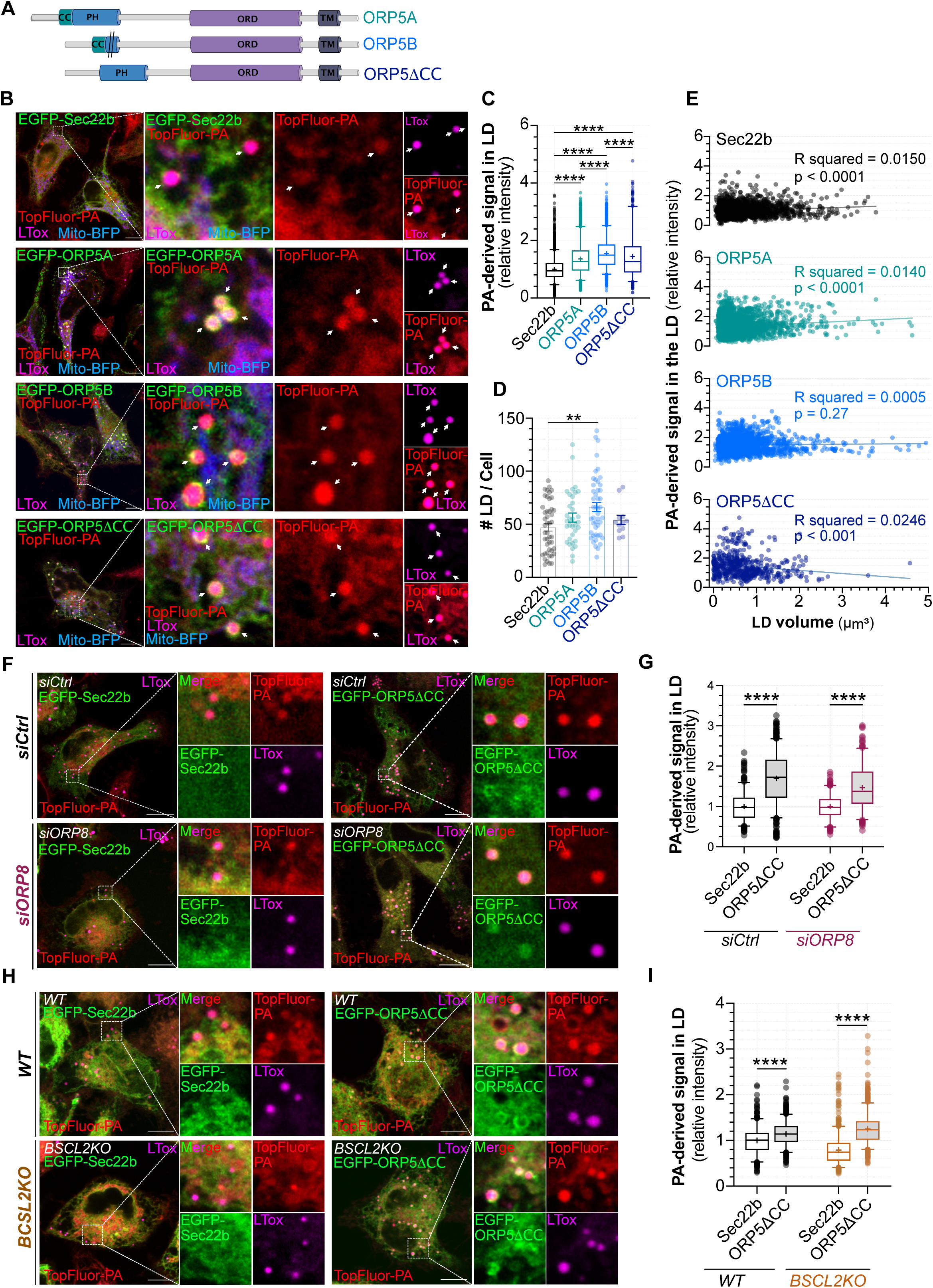
ORP5 facilitates the metabolism of phosphatidic acid into triacylglycerols and its storage in lipid droplets in a seipin/ORP8-independent-manner. **(A)** Graphic illustration of ORP5 domains. ORP5A represents the full length ORP5 containing a coiled coil (CC) domain, a Pleckstrin homology (PH) domain, an ORD domain (lipid transfer domain) and a transmembrane (TM) domain. ORP5B represents a natural isoform of ORP5 lacking the PH domain, and ORP5ΔCC represents an ORP5 mutant lacking the CC domain. **(B)** Representative confocal images, including full cell and zoomed regions, of HeLa cells co-transfected with Mito-BFP (blue), to mark mitochondria, and EGFP-Sec22b (green), EGFP-ORP5A (green), EGFP-ORP5B (green) or EGFP-ORP5ΔCC (green). Cells were treated with 2 µM TopFluor-TMR-PA (red) for 30min and stained with LipidTox (LTox, purple) to mark lipid droplets. Scale bar, 10 µm. **(C)** Relative quantification TopFluor-TMR-PA-derived fluorescence intensity co-localizing with lipid droplets. ORP5 facilitates the accumulation of TopFluor-TMR-PA-derived-signal in the lipid droplets, with EGFP-ORP5B having the strongest effect. Data are represented as boxplots showing the interquartile range, the median and the mean (+) of *n* = 577 - 3266 lipid droplets in at least three independent experiments. Statistical significance between data sets was assessed using Kruskal–Wallis test followed by multiple comparisons, with ****p < 0.0001. **(D)** Number of lipid droplets per cell expressing EGFP-Sec22b, EGFP-ORP5A, EGFP-ORP5B or EGFP-ORP5ΔCC. ORP5B induces a significant increase in the number of lipid droplets as compared to control cells expressing the general ER marker EGFP-Sec22b. Data are represented as mean ± SEM of *n* = 12 - 40 cells. Statistical significance between data sets was assessed using Kruskal–Wallis test followed by multiple comparisons, **p < 0.01. **(E)** Scatter plot of TopFluor-TMR-PA derived signal intensity and lipid droplets volume in cells expressing EGFP-Sec22b, EGFP-ORP5A, EGFP-ORP5B or EGFP-ORP5ΔCC. Data points, 577 - 3266 lipid droplets, were fitted with simple linear regression with P value and R-squared indicated in the plots. Top-Fluor-TMR-PA accumulation in the lipid droplets is independent of the lipid droplets size. **(F)** Representative confocal images, including full cell and zoomed regions, of control and ORP8-depleted HeLa cells expressing EGFP-Sec22b (green) or EGFP-ORP5ΔCC (green). Cells were treated with 2 µM TopFluor-TMR-PA (red) for 30min and lipid droplets were stained with LipidTox (purple). Scale bar, 10 µm. **(G)** Relative quantification TopFluor-TMR-PA-derived fluorescence intensity co-localizing with lipid droplets in control and ORP8-depleted cells expressing EGFPsec22b or EGFP-ORP5ΔCC. When compared with EGFP-Sec22b, EGFP-ORP5ΔCC promotes an increase of TopFluor-TMR-PA-derived-signal in the lipid droplets in both control and RNAi targeting ORP8-treated cells. Data are represented as boxplots showing the interquartile range, the median and the mean (+) of *n* = 244 - 589 lipid droplets. Statistical significance between data sets was assessed using Kruskal–Wallis test followed by multiple comparisons, with ****p < 0.0001. **(H)** Representative confocal images, full cell and zoomed regions, of *WT* and seipin knockout (*BCSL2KO)* HeLa cells expressing EGFP-Sec22b (green) or EGFP-ORP5ΔCC (green). Cells were treated with 2 µM TopFluor-TMR-PA (red) for 30min and lipid droplets were stained with LipidTox (LTox, purple). Scale bar, 10 µm. **(I)** Relative quantification TopFluor-TMR-PA-derived fluorescence intensity co-localizing with lipid droplets in *WT* and *BCSL2KO* cells expressing EGFPsec22b or EGFP-ORP5ΔCC. EGFP-ORP5ΔCC promotes an increase of TopFluor-TMR-PA-derived-signal in the lipid droplets in both *WT* and *BCSL2KO* cells as compared with EGFP-Sec22b condition. Data are represented as boxplots showing the interquartile range, the median and the mean (+) of *n* = 501 - 809 lipid droplets. Statistical significance between data sets was assessed using Kruskal–Wallis test followed by multiple comparisons, with ****p < 0.0001.

Internalized TopFluor-TMR-PA displayed a broad distribution at the PM, ER, mitochondria, and LDs, as revealed by confocal and structured illumination microscopy (SIM) (Fig. 2B, Supp. Fig. 5). Notably, TopFluor-TMR-PA-derived signal accumulated in LDs of cells expressing ORP5 isoforms (Fig. 2B). Quantification by 3D IMARIS analysis showed significantly increased LD-localized TopFluor-TMR-PA fluorescence in cells expressing EGFP-ORP5A, ORP5B, or ORP5ΔCC compared with EGFP-Sec22b control (Fig. 2C), with ORP5B exerting the strongest effect (Fig. 2C). Consistently, ORP5B overexpression also increased LD number per cell, whereas ORP5A and ORP5ΔCC had more moderate effects (Fig. 2D). Linear regression analysis further revealed that the incorporation of PA-derived fluorescence into LDs was independent of LD size across all conditions, as indicated by low R squared values (Fig. 2E). Together, these data confirm that ORP5 promotes the metabolism of PA into TAG and its storage in LDs.

Since ORP5 and ORP8 localize as a complex at multiple MCS, including Mito–ER–LD junctions (Chung et al. 2015; Monteiro-Cardoso et al. 2022; Guyard et al. 2022), and share similar domain structure and overlapping functions, we asked whether ORP8 also contributes to PA metabolism. Strikingly, depletion of ORP8 did not alter LD-associated TopFluor-TMR-PA– derived fluorescence (Suppl. Fig. 6A,B), indicating that, unlike ORP5, ORP8 does not facilitate PA incorporation into TAG for LD storage.

We next sought to provide further evidences that ORP5 function in directing PA towards LDs is independent of ORP8. To this end, we transfected control and ORP8-depleted HeLa cells with EGFP-Sec22b or EGFP-ORP5ΔCC, a mutant lacking the coiled-coil domain required for ORP8 binding. Since the expression of ORP8 was not completely abolished by siRNA treatment (Supp. Fig 6C), we used ORP5ΔCC to circumvent residual interaction with endogenous ORP8. As expected, EGFP-ORP5ΔCC expression in control cells (*siCtrl*) significantly increased TopFluor-TMR-PA–derived fluorescence in LDs compared with EGFP-Sec22b–expressing cells, and this effect was maintained in ORP8-depleted cells (Fig. 2F, G). These results demonstrate that ORP5 regulates PA storage in LDs independently of ORP8.

We have previously shown that ORP5 controls seipin targeting to MAM–LD contacts (Guyard et al. 2022), and seipin oligomers have been reported to trap PA and DAG in yeast (Zoni et al. 2021; House et al. 2025; Yan et al. 2018; Choudhary et al. 2020). Therefore, we then tested whether seipin is required for ORP5-mediated PA flux into LDs. For that we expressed EGFP-ORP5ΔCC in seipin (*BSCL2*) knockout (*BSCL2KO*) HeLa cells and monitored TopFluor-TMR-PA–derived LD signal. In agreement with prior and recent reports implicating seipin in PA handling and LD formation (Salo et al. 2019; Yan et al. 2018; Romanauska et al. 2024; House et al. 2025), seipin-deficient cells, either untransfected or transfected with EGFP-Sec22b, displayed markedly reduced PA incorporation into LDs (Suppl. Fig. 6D,E; Fig. 2H, I). Strikingly, expression of EGFP-ORP5ΔCC rescued this phenotype, restoring PA incorporation into LDs (Fig. 2H, I). These findings indicate that ORP5 can direct PA toward LD storage independently of seipin.

### ORP5 enriches PA at MAM and regulates its distribution across mitochondria and lipid droplets

We next tested whether ORP5 activity leads to PA conversion into DAG via Lipin-1. Cells expressing EGFP-ORP5B were treated with propranolol, a Lipin-1! inhibitor. In untreated cells, ORP5B expression promoted strong accumulation of TopFluor-TMR-PA-derived signal in the LD core (Fig. 3A,B). In contrast, propranolol treatment abolished PA incorporation into LDs, and instead PA-derived fluorescence concentrated in ER subdomains positive for EGFP-ORP5B, tightly wrapping LDs (Fig. 3A,B). This effect was not observed in Sec22b-expressing cells (Fig. 3A,B), indicating a specific role for ORP5. Consistent with our previous findings that ORP5 localizes at MAM-LD contacts, these ORP5- and PA-positive ER-subdomains wrapping LDs were closely associated with mitochondria (Fig. 3A).

**Figure 3:**
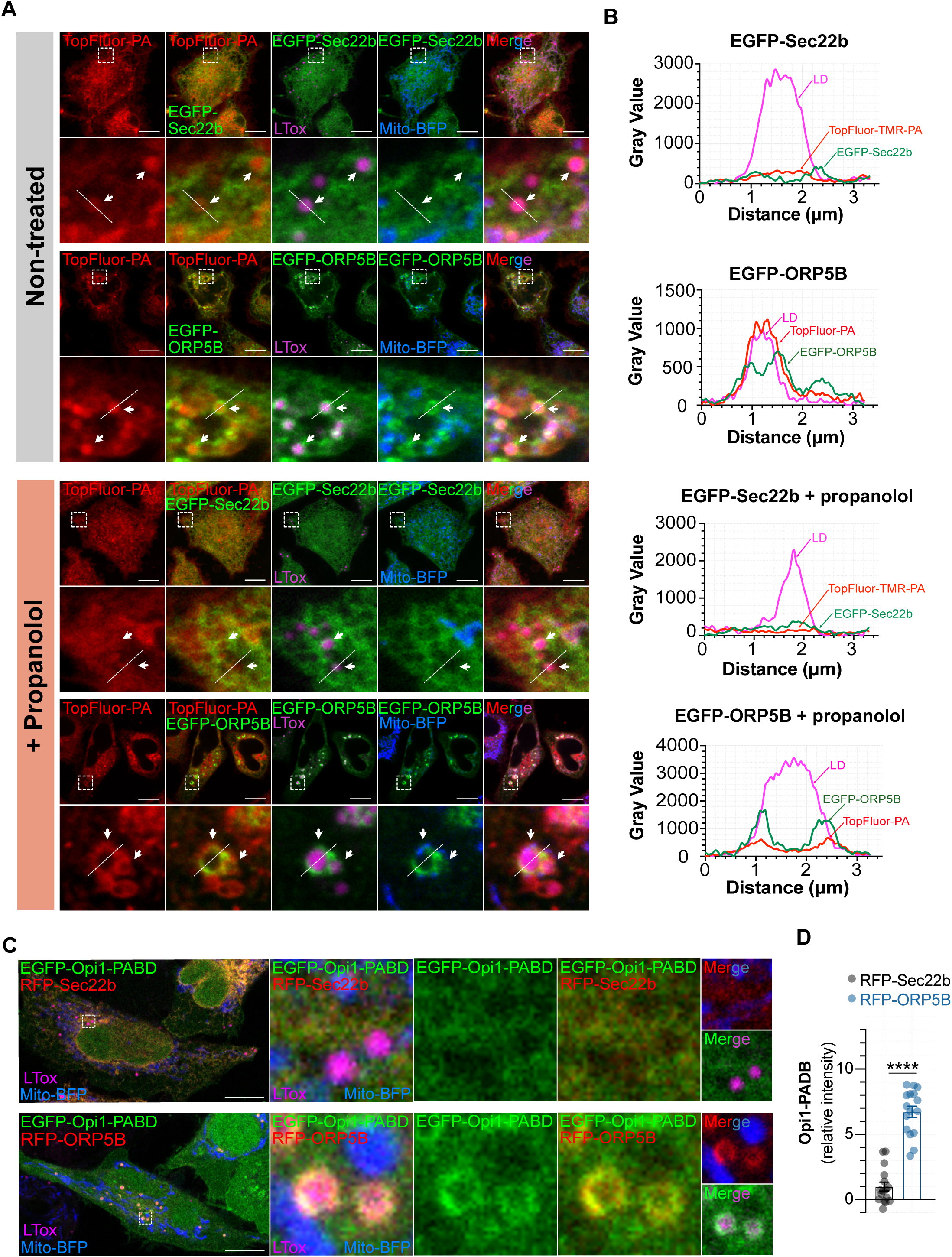
ORP5 stabilizes phosphatidic acid at MAM-lipid droplet interface. **(A)** Representative confocal images of zoomed regions of HeLa cells expressing EGFP-Sec22b (green) or EGFP-ORP5B (green) and Mito-BFP (blue) and stained with LipidTox to mark lipid droplets (LTox, purple). Cells were treated with 2 µM TopFluor-TMR-PA for 30 min (upper panel) or with 100µM propranolol (Lipin-1! inhibitor) for 1h plus 2 µM TopFluor-TMR-PA for 30min (lower panel). Scale bar,10 µm. Dotted lines represent the regions plotted in fluorescence intensity profiles. **(B)** Fluorescence intensity profiles corresponding to the intensities of EGFP-Sec22b(green) or EGFP-ORP5B (green), TopFluor-TMR-PA (red) and lipid droplet (purple) in cells non-treated or treated with propranolol, as indicated in the figure. Propranolol blocks the conversion of PA in DAG. While in non-treated cells TopFluor-TMR-PA accumulates in the lipid droplets, in propranolol-treated cells it accumulates where EGFP-ORP5B localizes, particularly in region where ORP5 is in close association with mitochondria and wrapping the lipid droplets. **(C)** Illustrative confocal images, whole cells and zoomed regions, of HeLa cells co-expressing EGFP-OPi1-PABD (PA sensor, green) and Mito-BFP (blue) with either RFP-Sec22b (red) or RFP-ORP5B (red). Lipid droplets were stained with LipidTox (LTox, purple). Scale bar, 10 µm. **(D)** Relative quantification of EGFP-OPi1-PABD fluorescence intensity surrounding the Lipid droplets. EGFP-OPi1-PABD fluorescence intensity in close association with lipid droplets is increased in cells expressing RFP-ORP5B as compared to cells expressing RFP-Sec22b. Data are represented as mean ± SEM of *n* = 16 lipid droplets. Statistical significance between data sets was assessed using Mann-Whitney U test, ****p < 0.0001.

Since PA also serves as a precursor for multiple phospholipids (Zhou et al. 2024), we next examined whether ORP5 recruits or stabilizes endogenous PA using the sensor EGFP-Opi1-PABD (PA-binding domain). In Sec22b-expressing cells, Opi1-PABD displayed diffuse cytoplasmic localization (Fig. 3C). In contrast, ORP5B expression led to strong Opi1-PABD accumulation at ER subdomains where ORP5B wrapped LDs, often adjacent to mitochondria (Fig. 3C,D). Together, these results indicate that ORP5 recruits and stabilizes PA at MAM–LD contact sites, promoting its conversion into TAG and incorporation into LDs.

In recent years, LD-mitochondria crosstalk has emerged as a key metabolic axis (Freyre et al. 2019; Monteiro-Cardoso and Giordano 2024), with PA playing a central role in both organelles, as it serves as a metabolic intermediate for both TAG and cardiolipin (Fig. 4A). Because PA cannot be synthesized within mitochondria, it must be imported from the ER through MCS and the action of LTPs. While this ER-to-mitochondria transfer is well established, whether mitochondrial PA can be rerouted back to the ER and converted into TAG remains unclear. However, recent studies suggest that perturbations in mitochondrial PA handling can influence ER lipid metabolism (Eiyama et al. 2021).

**Figure 4:**
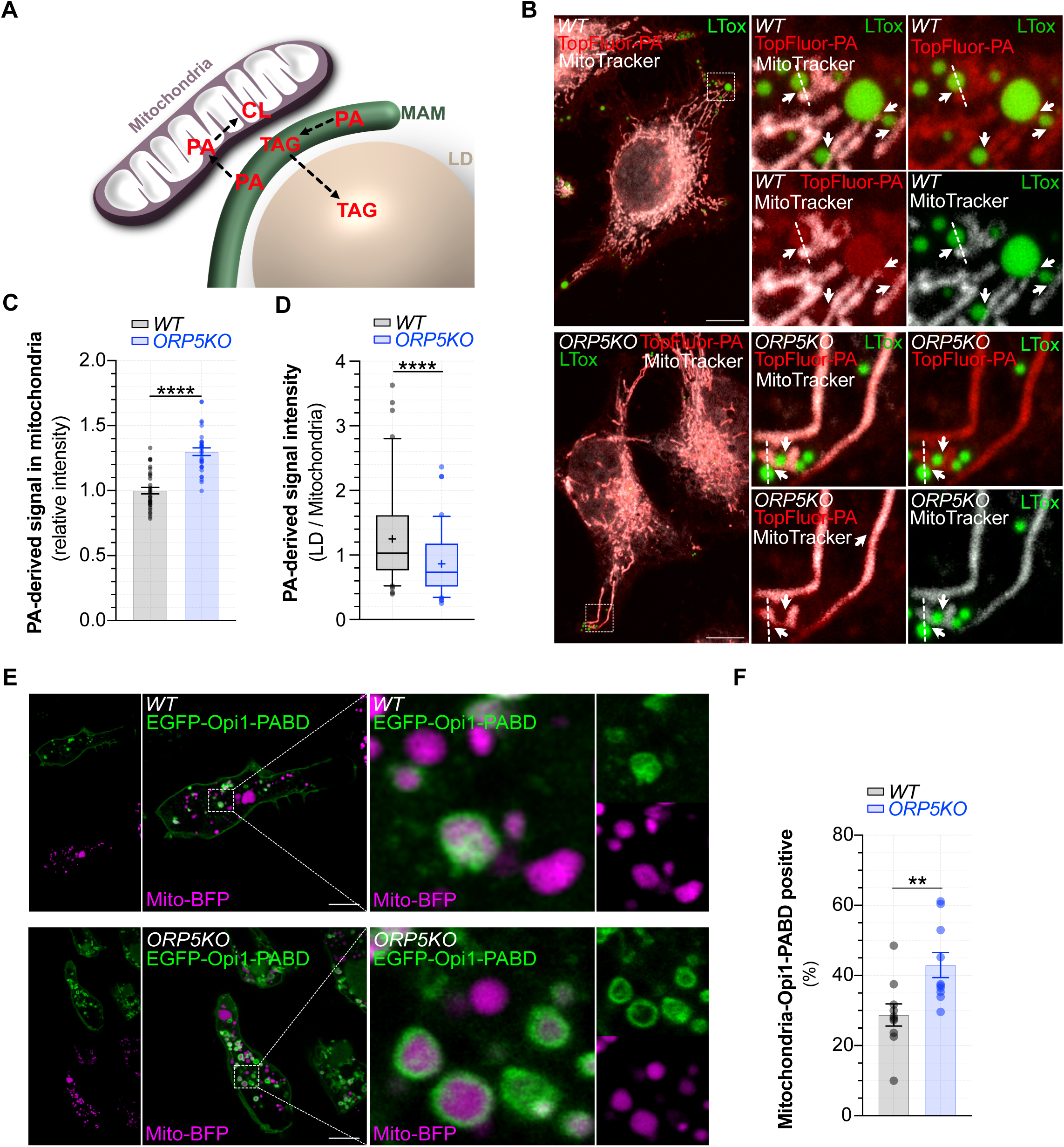
ORP5 maintains the balance of phosphatidic acid between storage in lipid droplets and incorporation in mitochondrial membranes. **(A)** Schematic representation of phosphatidic acid (PA) metabolism across mitochondria-associated endoplasmic reticulum (ER) membranes - lipid droplet (LD) interface. In the ER, PA is converted into triacylglycerols (TAG) to be stored in lipid droplets (LD). Additionally, ER-derived PA is shuttled to the mitochondria where it is converted into cardiolipin (CL). **(B)** Illustrative confocal images, full cell and zoomed regions, of *WT* and *ORP5KO* HeLa cells treated with 2 µM TopFluor-TMR-PA (red) for 30min. Mitochondria and lipid droplets were stained with MitoTracker deep red (gray) and LipidTox (purple), respectively. Scale bar, 10 µm. **(C)** Relative quantification of TopFluor-TMR-PA intensity associated to mitochondria. Mitochondria in *ORP5KO* cells display increased TopFluor-TMR-PA. Data are represented as mean ± SEM of *n* = 32 cells. Statistical significance between data sets was assessed using Mann-Whitney U test, ****p < 0.0001. **(D)** LD to mitochondria TopFluor-TMR-PA-derived fluorescence intensity ratio in *WT* and *ORP5KO* cells. Depletion of ORP5 results in a significant decrease in LD / mitochondria TopFluor-TMR-PA-derived signal. Data are represented as boxplots showing the interquartile range, the median and the mean (+) of *n* = 91 - 98. Statistical significance between data sets was assessed using Mann-Whitney U test, with ****p < 0.0001. **(E)** Confocal images showing whole cells and zoomed areas of swelled HeLa *WT* and *ORP5KO* cells expressing EGFP-OPi1-PABD (green), where mitochondria have been marked with Mito-BFP (purple). **(F)** Percentage of mitochondrial vesicles marked by EGFP-OPi1-PABD. ORP5-deleted cells display increased EGFP-OPi1-PABD-posite mitochondrial vesicles. Data are represented as mean ± SEM of *n* = 10 cells. Statistical significance between data sets was assessed using Mann-Whitney U test, **p < 0.01.

We hypothesized that ORP5 maintains PA balance between LDs and mitochondria. To test this, we incubated *W*T and ORP5-depleted (*ORP5KO* and *siORP5*) cells with TopFluor-TMR-PA, and visualized LDs and mitochondria with LipidTox and either Mitotracker or transfection of Tom20-StayGold, respectively. Confocal microscopy analysis revealed that in *WT* cells, PA was efficiently directed to LDs, whereas in ORP5-deficient cells, it accumulated more prominently in mitochondria, showing greater colocalization with Mitotracker (*ORP5KO*) or Tom20-StayGold (*siORP5*) than *WT* or *siCtrl* cells (Fig. 4B, Supp. Fig. 3B). Quantification confirmed that PA-derived fluorescence signal was increased in the mitochondria of *ORP5KO* cells (Fig. 4C). Additionally, a reduced LD-to-mitochondria PA ratio was observed in *ORP5KO* cells (Fig. 4D), consistent with a redistribution of PA toward mitochondria.

To validate these findings with an independent approach, we expressed the PA biosensor Opi1-PABD in *WT* and *ORP5KO* cells. Following hypotonic swelling to improve its visualization (Santinho et al. 2024; Santinho et al. 2020), we observed, in both *WT* and *ORP5KO* cells, two distinct mitochondria populations, one enriched in Opi1-PABD, and another lacking Opi1-PABD enrichment (Fig. 4E). We assessed the % of each one of these population and, interestingly, we found an increase of Opi1-PABD-positive mitochondria in cells depleted of ORP5 (Fig. 4F). These results indicate elevated PA levels at the outer mitochondrial membrane in the absence of ORP5 and support a model in which ORP5 stabilizes PA at MAM and maintains its partitioning between mitochondria and LDs, thereby preventing its excessive accumulation in mitochondrial membranes.

### ORP5 regulates the levels of specific cardiolipin and triacylglycerol species

The redistribution of PA in ORP5-deficient cells prompted us to examine how this affects mitochondrial and LD lipid composition. To this end, we isolated MAM and mitochondrial fractions from ORP5-depleted and control cells using differential centrifugation combined with a Percoll density gradient, as described previously (Monteiro-Cardoso et al. 2022; Galmes et al. 2016), and performed lipidomic analysis (Fig. 5 A; Supp. Fig. 7). Lipid extracts were isolated from three different MAM and mitochondria preparations and analyzed by mass spectrometry. We detected two PA species, 32:0 (16:0 / 16:0) and 34:1 (16:0 / 18:1), in the lipid extract from both ORP5-depleted (*siORP5*) and control (*siCtrl)* MAM. Interestingly, the levels of both PA 32:0 and 34:1 were slightly decreased in the absence of ORP5 (Fig. 5B), consistent with impaired PA stabilization at MAM. In mitochondria, PA levels were below detection due to rapid conversion into cardiolipin. Notably, however, ORP5 depletion caused a significant increase in several cardiolipin species, particularly cardiolipins 68:4 and 68:5 (Fig. 5C). This supports the idea that excess mitochondrial PA is funneled into cardiolipin synthesis when ORP5 is absent.

**Figure 5:**
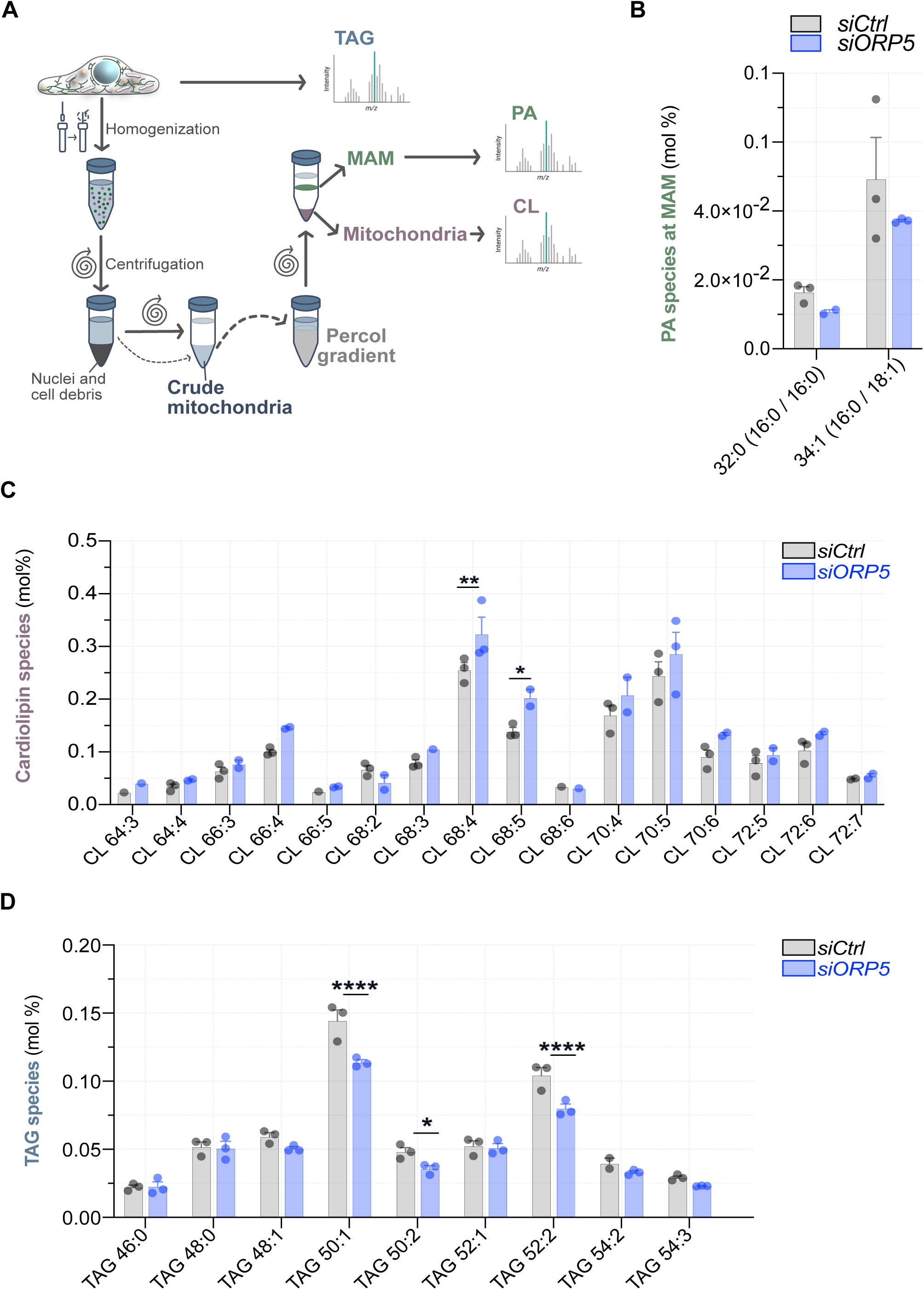
ORP5 affects the levels of specific cardiolipin and triacylglycerol species. **(A)** Schematic representation of conceptual framework of mitochondria and mitochondria-associated ER-membranes (MAM) isolation from controls and ORP5-depleted HeLa cells for mass spectrometry analysis. **(B)** Phosphatidic acid (PA) content in MAM isolated from control HeLa cells (*siCtrl*) and RNAi treated HeLa cells targeting ORP5 (*siORP5*)**. (C)** Content of cardiolipin (CL) species in mitochondrial membranes of control (*siCtrl*) and ORP5-depleted (siORP5) HeLa cells. Absence of ORP5 results in increased levels of cardiolipin 68:4 and 68:5. **(D)** Content of triacylglycerols (TAG) species in total control (*siCtrl*) and ORP5-depleted (*siORP5*) HeLa cells. Depletion of ORP5 decreased the levels of TAG 50:1, 50:2 and 52:2. Data are presented as the mean of mol% ± SEM of *n* = 3 independent experiments. *P* values were calculated using unpaired student’*t* test. *p < 0.05; **p < 0.01; ****p < 0.0001. Cardiolipin and TAG species are represented by the total number of carbon atoms (first number) and by the total number of double bonds (second number).

Given our imaging results and the observed decrease in bulk TAG upon ORP5 loss, we also profiled TAG species in lipid extracts isolated from *siORP5* and *siCtrl* HeLa cells by mass spectrometry. ORP5 depletion selectively reduced TAG 50:1, 50:2, and 52:2 (Fig. 5D). Thus, ORP5 regulates PA metabolism at MAM by stabilizing specific PA species, which in turn influences both cardiolipin synthesis in mitochondria and the production of defined TAG species in LDs. Whether ORP5 also regulates the transfer of PA at the MAM-LD junction remains to be determined.

### ORP5 transfers PA *in vitro* and favors its transport from the mitochondria to the ER

The ORD domain of ORP5 has been shown to transfer the anionic phospholipid PS between liposomes, either in counter transport with PI(4)P and PI(4,5)P_2_, or independently of other lipid gradients (Chung et al. 2015; Ghai et al. 2017; Monteiro-Cardoso et al. 2022). In contrast, it does not transport the zwitterionic phospholipids phosphatidylcholine (PC) and phosphatidylethanolamine (PE) *in vitro* (Monteiro-Cardoso et al, 2022). In cells, ORP5 mediates the transport of PS at ER-PM contacts in exchange with PM PI(4)P or PI(4,5)P_2_ (Ghai et al. 2017; Chung et al. 2015), and at ER-mitochondria contacts unidirectionally, driven by the PS concentration gradient (Monteiro-Cardoso et al. 2022). However, whether the ORD domain of ORP5 can also transfer other lipids has remained unclear.

Since PA, like PS, is an anionic phospholipid, we next tested whether ORP5 could also transfer PA. To this end, we purified the recombinant ORP5-ORD (ORD5) from bacteria (Suppl. Fig. 8A) and assessed its ability to transport fluorescent phospholipids (Top-Fluor PS, TopFluor-TMR-PA, and Top-Fluor PE) from donor to acceptor liposomes using a previously established *in vitro* lipid transport assay (Monteiro-Cardoso et al. 2022). Donor liposomes contained fluorescent PA (TopFluor-TMR-PA, 18:1/6:0; oleoyl at one acyl chain and a hexanoyl chain bearing a terminal BODIPY-TMR) or other fluorescent lipids, together with biotinylated lipids for immobilization on streptavidin beads. Immobilized donor liposomes were then incubated with acceptor liposomes composed of either PC:PE or PC alone. After 1 h at 37 °C, acceptor liposomes were recovered from the supernatant and their fluorescence was measured (Fig. 6A; Suppl. Fig. 8B).

**Figure 6:**
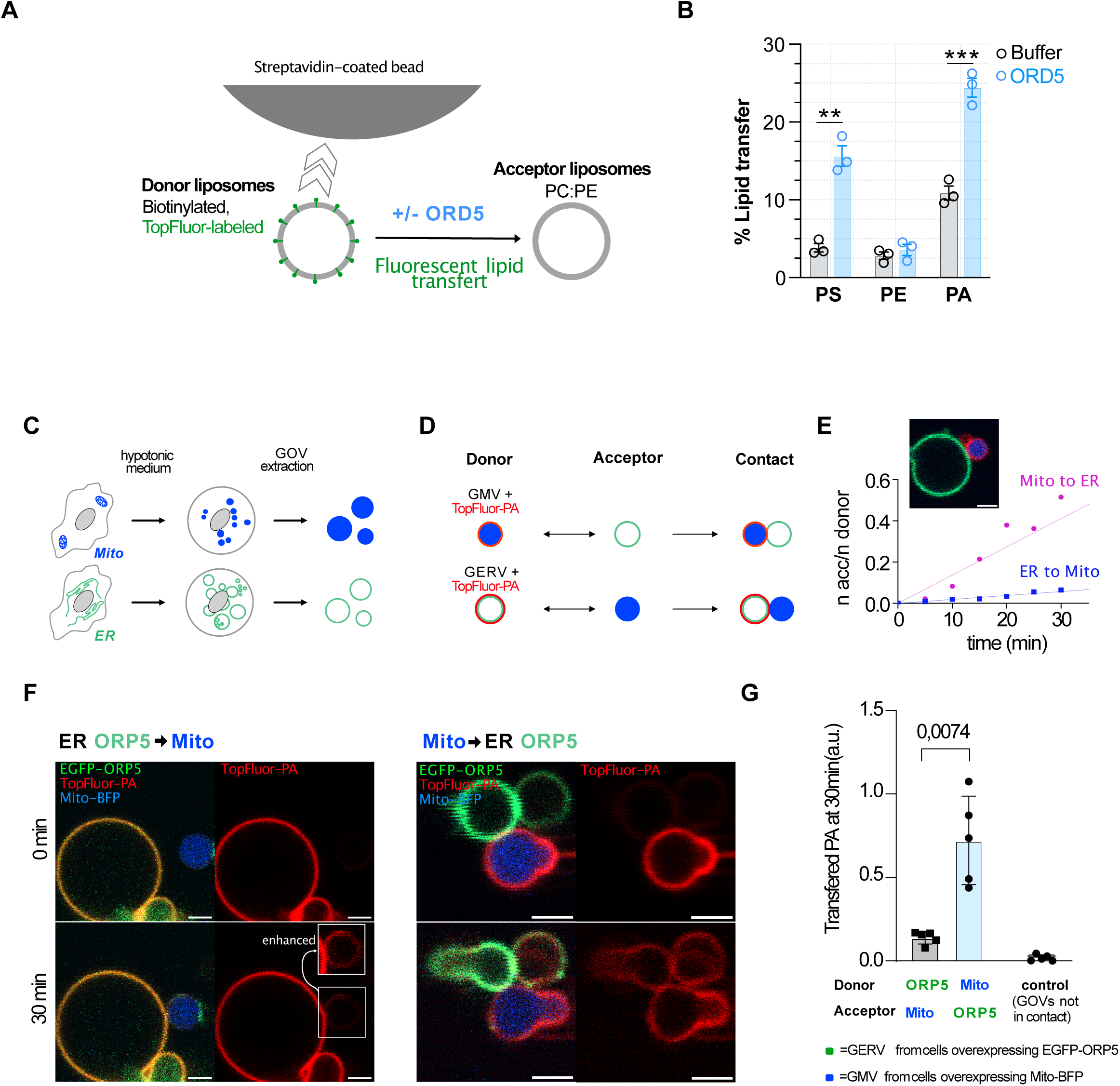
ORP5 mediates the transfer of phosphatidic acid *in vitro* between liposomes and *ex vivo* between giant ER (GERV) and mitochondria (GMitoV) vesicles. (**A**) Schematic representation of the conceptual framework of *in vitro* lipid transfer assay between liposomes mediated by the ORD domain of ORP5 (ORD 5). Donor Liposomes contain POPC, Biotinalited and TopFluor phospholipids. Acceptor Liposomes are composed of a mixture of POPC and Egg trans phosphatidylethanolamine (PE). **(B)** Percentage of transfer of TopFluor-phospholipid phosphatidylserine (PS), PE and phosphatidic acid (PA), mediated by ORD5 between the donor and the acceptor liposomes. Data are presented as % of transferred lipid ± SEM and are the mean of three independent experiments. Statistical analysis were performed using unpaired student’s *t*-test, **p <0.01, ***p<0.001. **(C)** Schematic representation of conceptual framework followed to investigate ORP5-mediated PA transfer using giant organelles vesicles (GOVs). HeLa cells transfected with either ER EGFP-ORP5B (green) or Mito-BFP (blue) were exposed to hypotonic media, and GOVs were extracted. **(D)** Donor GOVs (either giant ER vesicles (GERV) or giant Mitochondria vesicles (GMV)) fed with TopFluor-TMR-PA (red) for 30min, combined with acceptor GOVs and brought into contact via micropipettes. **(E)** Representative relative PA fluorescence intensity increases over time after established contact of donor and acceptor GOVs with linear regression fit. **(F)** PA transfer between GOVs upon artificially established contact, comparing representative confocal images at 0 min (top) and after 30 min (bottom) for the donor to acceptor pairs: ER ORP5 to Mito (left) and Mito to ER ORP5 (right), with TopFluor-TMR-PA (red) transferring from donor to acceptor GOVs. Inlet in TopFluor-TMR-PA channel 30 min for condition ER ORP5◊Mito shows enhanced brightness. **(G)** Quantification of the share of transferred PA from donor to acceptor organelle after 30 min for various donor-acceptor pairs. Depicting a faster transfer rate for a transfer from Mito to ER. For the control the Mito to ER ORP5 condition was observed over time with no artificial contact. Sample preparation was done at least 3 different times. For all Welch t-test was used to analyze statistical differences.

As expected, ORD5 efficiently transferred PS but not PE between liposomes *in vitro* (Fig. 6B, Suppl. Fig. 8C). Remarkably, we also observed robust PA transfer, demonstrating that in addition to PS, the ORD domain of ORP5 can mediate PA transport *in vitro* (Fig. 6B; Suppl. Fig.8C). This finding suggests that ORP5 may facilitate PA transport between organelles in cells.

We have previously shown that ORP5 predominantly localizes at ER-mitochondria contact sites where it mediates transfer of PS (Monteiro-Cardoso et al. 2022), and regulates LDs biogenesis (Guyard et al. 2022). We investigated whether ORP5 can mediate non-vesicular transport of PA between mitochondria and the ER. To this end, we developed an *ex vivo* lipid transfer assay based on lipid exchange between distinct giant organelle vesicles (GOVs), namely giant ER vesicles (GERVs) and giant mitochondrial vesicles (GMVs) (Santinho et al. 2024). Cells were transfected with either EGFP–ORP5B or Mito–BFP to label luminal mitochondria, and then subjected to a hypotonic medium. The following plasma membrane disruption (Santinho et al. 2024), enabled the extraction of EGFP–ORP5B–positive GERVs and Mito–BFP–positive GMVs in separate dishes (Fig. 6C). Either GERVs or GMVs were then selectively loaded with TopFluor–TMR–PA, allowing us to designate GMVs or GERVs as the PA donor GOV and the reciprocal GOV as the acceptor (Fig. 6D). Pairs of different GOVs were subsequently brought into contact for 40 min (Supp. Fig. 8D), using a micropipette aspiration setup. Lipid transfer was detected as a time-dependent, specific increase in PA fluorescence in the acceptor GOV (Fig. 6E). Strikingly and reproducibly, PA transport was consistently much higher from mitochondria to the ER than in the reverse direction (Fig. 6F– G). These results support a preferential and favorable transport of PA from mitochondria to the ER at MAMs.

Together, these *in vitro* and *ex vivo* assays suggest that ORP5 can transfer PA via its ORD domain and favor its transport from mitochondria to the ER.

### ORP5 regulates mitochondria morphology

Mitochondria continuously remodel their network through opposing processes of fission and fusion, collectively referred to as mitochondrial dynamics (Tilokani et al. 2018). These events are essential to maintain mitochondrial function and to adapt to cellular energy demands. Mitochondrial dynamics are tightly regulated by protein machineries and by lipid composition, including PA and cardiolipin (Kameoka et al. 2018). Both lipids directly interact with key fusion and fission proteins—mitofusins 1/2 (MFN1/2), OPA1, and DRP1—and lipid-modifying enzymes that alter PA or cardiolipin levels have been shown to influence the balance between fission and fusion, impacting mitochondrial morphology (Kameoka et al. 2018; Tilokani et al. 2018).

Because ORP5 deficiency perturbs PA distribution at MAM–LD contacts and increases cardiolipin levels in mitochondria, we hypothesized that ORP5 loss might impact mitochondrial dynamics and morphology. To test this, we performed morphometric analysis of mitochondria stained with MitoTracker in control (*siCtrl*) and ORP5-depleted (*siORP5*) cells. Remarkably, *ORP5KD* cells displayed significantly elongated mitochondria compared with controls, as reflected by increased aspect ratio (AR; Fig. 7A,B). Form factor (FF) analysis further revealed enhanced mitochondrial branching and interconnectivity (Fig. 7A,C), accompanied by a reduction in mitochondrial number per cell (Fig. 7D).

**Figure 7:**
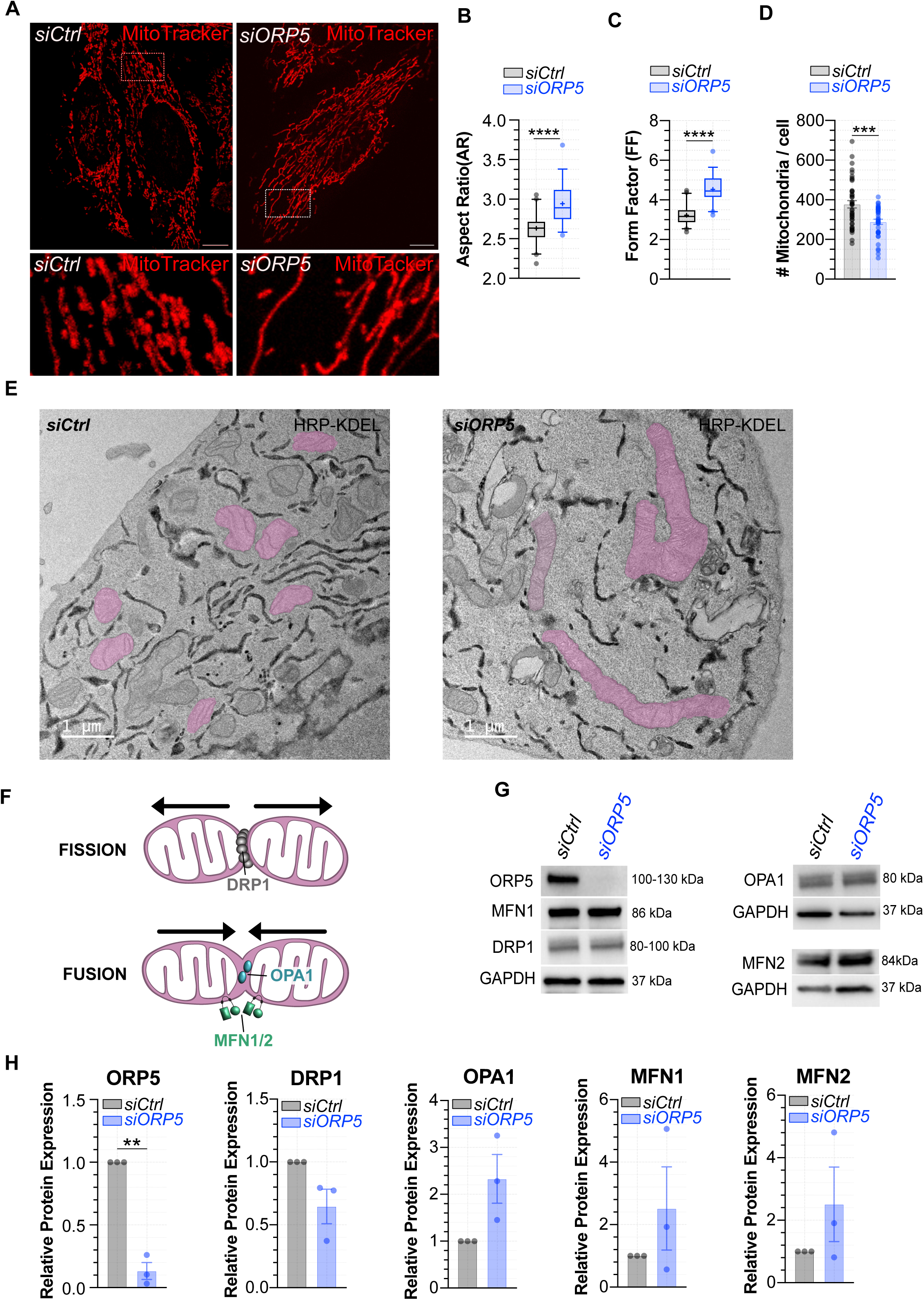
ORP5 alters mitochondria morphology. **(A)** Representative confocal images, including full cell and zoomed regions, showing mitochondria morphology in control (*siCtrl*) and RNAi ORP5-depleted (*siORP5*) HeLa cells stained with mitotracker orange (red) to mark mitochondria. **(B-D)** Quantification of the mitochondrial shape descriptors aspect ratio (AR), indicator of mitochondria elongation, and form factor (FF), indicator of mitochondrial branching, and the number of mitochondria per cell. *siORP5* cells display less and more elongated mitochondria as compared with *siCtrl* cells. Data are represented as boxplots showing the interquartile range, the median and the mean (+) of *n* = 39 - 42 cells in three independent experiments. Statistical significance between data sets was assessed using Mann-Whitney U test, ***p < 0.001, ****p < 0.0001. **(E)** Electron micrograph of ultrathin cryosections of control (*siCtrl*) and ORP5-depleted (*siORP5*) HeLa cells, showing elongated mitochondria in *siORP5* HeLa cells. **(F)** Simplified schematic representation of mitochondria fission and fusion and the core protein involved in these processes, dynamin-related protein 1 (DRP1, fission), Mitofusins 1/2 (MFN1/2, fusion) and OPA1 mitochondrial dynamin like GTPase (fusion). **(G)** Representative immunoblot showing ORP5, DRP1, MFN1, MFN2, and OPA1 protein levels in HeLa cells 48 h after transfection with control siRNA or with a *ORP5*-specific siRNA. **(H)** Relative Protein levels ORP5, DRP1, MFN1, MFN2, and OPA1 in control and ORP5-depleted cells. Data are presented as the mean ± SEM of three independent experiments. *P* values were calculated using unpaired student’*t* test. **p < 0.01.

To assess whether these effects were specific to ORP5, we depleted its binding partner ORP8, which also mediates PS transfer at ER–mitochondria contacts (Monteiro-Cardoso et al. 2022). Unlike ORP5, ORP8 depletion did not alter mitochondrial AR or FF (Supp. Fig. 9A– C), indicating that the mitochondrial hyperfused phenotype is uniquely dependent on ORP5 loss of activity. Notably, the elongated and hyperconnected mitochondrial phenotype observed by confocal microscopy in ORP5-deficient cells was also evident at the ultrastructural level by transmission electron microscopy (Fig. 7E).

We next examined whether permanent loss of ORP5 yields comparable changes. Consistent with the knockdown results, *ORP5KO* cells exhibited significantly elongated and branched mitochondria relative to *WT* cells, as indicated by increased AR and FF values (Suppl. Fig. 9E–G). These findings suggest that ORP5 deficiency shifts mitochondrial dynamics toward fusion and/or reduces fission, resulting in fewer but more interconnected mitochondria.

To determine whether these morphological changes correlate with altered expression of mitochondrial dynamics proteins, we assessed DRP1, OPA1, and MFN1/2 by immunoblotting. Although ORP5 depletion modestly decreased DRP1 and slightly increased OPA1 and MFN1/2 levels, none of these changes reached statistical significance (Fig. 7F–H). This suggests that the hyperfused mitochondrial phenotype does not arise from altered expression of the core fission–fusion machinery. Instead, our results indicate that ORP5 regulates mitochondrial morphology primarily through remodeling of mitochondrial lipid composition, particularly by controlling the levels and distribution of PA and cardiolipin.

## DISCUSSION

Our study identifies ORP5 as a critical regulator of PA partitioning across mitochondria, ER and LD at the tripartite MAM-LD junction. By integrating imaging, biochemical, and lipidomic approaches across *in situ*, *in vitro* and *ex vivo* assays, we demonstrate that ORP5 promotes the efficient channeling of PA into TAG and LD storage, while simultaneously preventing its excessive accumulation in mitochondria, and subsequent cardiolipin overload (Supp. Fig. 9G). *In vitro* liposome assays and *ex vivo* experiments on isolated giant organelles from swollen cells revealed that ORP5 has a PA transfer activity and preferentially transfers PA from mitochondria toward the ER rather than the reverse (Fig. 6; Suppl. Fig. 5 D-E and 8B), highlighting its capacity to couple mitochondrial PA pools to LD biogenesis. ORP5 stabilizes PA at MAM subdomains (Fig. 3), facilitates its transfer across mitochondria–ER–LD interfaces (Fig. 6), and thereby maintains the balance between lipid storage and mitochondrial lipid biosynthesis. Loss of ORP5 disrupts this balance, resulting in specific TAG species deficits (Fig. 1, Fig. 5D), decrease in PA levels at MAMs (Fig. 4), increased levels of cardiolipin (Fig. 5 C), and mitochondrial hyperfusion (Fig. 7), revealing ORP5 as a previously unrecognized PA transfer protein that couples LD biogenesis to mitochondrial lipid metabolism and dynamics.

Previous work established ORP5 (and ORP8) as a PS transfer protein at ER–plasma membrane and ER-LD contacts, operating via counter-exchange with phosphoinositides, such as PI(4)P and/or PI(4,5)P_2_ (Chung et al. 2015; Ghai et al. 2017; Du et al. 2020). ORP5 and ORP8 can also transfer PS at ER–mitochondria contact sites independently of PI(4)P (Monteiro-Cardoso et al. 2022; Galmes et al. 2016). However, whether ORP5 also acts on other physiological lipid substrates remained unclear. Here, we find that ORP5’s role in LD formation is independent of its PS transport function: a PS-binding mutant fully rescues the LD defect (Supp. Fig. 1), whereas ORP8 deletion has no effect on mitochondrial morphology or PA storage (Fig. 2F,G; Supp. Fig. 9A–C). These results suggest ORP5 has uniquely evolved to handle PA at contact sites, separate from its PS-related functions and independent of ORP8.

We previously showed that ORP5 recruits seipin at MAM-LD contacts, where seipin plays a crucial role in LD nucleation and PA sensing (Yan et al. 2018; Guyard et al. 2022; House et al. 2025). Here we show that ORP5 PA-handling function at MAM-LD contacts does not depend on seipin (Fig. 2H, I), revealing that ORP5 defines a novel and independent PA regulation pathway. Nonetheless, ORP5 may aid in seipin recruitment by enriching local PA concentrations at MAM subdomains, suggesting complementary roles in spatial lipid organization at LD formation sites.

PA is a precursor for mitochondrial cardiolipin and must be imported from the ER. The LTPs mediating this transport remain largely undefined, though candidates such as PTPIP51, ESyt1, and VPS13 have been implicated in ER–mitochondria lipid exchange (Yeo et al. 2021; Sassano et al. 2023; Janer et al. 2024; Kumar et al. 2018; Guillen-Samander et al. 2021). PTPIP51 has been suggested to facilitate PA transfer from ER to mitochondria, in addition to its tethering function, but its direct mechanistic role remains unclear (Yeo et al. 2021; Giordano 2021). While the direction from ER to mitochondria is established, whether mitochondrial PA can be rerouted back to the ER and converted into TAG remains an open question. Recent studies suggest that perturbations in mitochondrial PA handling can influence ER lipid metabolism (Eiyama et al. 2021), raising the possibility of bidirectional PA flux across MAMs. Our *in situ, in vitro,* and *ex vivo* data suggest that ORP5, and potentially other yet-to-be-identified LTPs, preferentially channel PA from mitochondria to the ER-LD axis, providing mechanistic evidence that mitochondrial PA pools can be mobilized for lipid storage and reinforcing the concept of dynamic, bidirectional lipid exchange at MAM–LD junctions.

Beyond their role in fatty acid oxidation, subsets of mitochondria adjacent to LDs can adopt an anabolic role, supporting lipid synthesis and storage (Benador et al. 2018; Klemm 2021; Freyre et al. 2019). These peridroplet mitochondria (PDM) are physically tethered to LDs and display distinct proteomic and metabolic features that promote TAG accumulation, distinguishing them from cytoplasmic mitochondria specialized for fatty acid catabolism (Najt et al. 2023). LD-associated mitochondria are therefore positioned to adjust their function depending on cellular metabolic state. MAMs serve as metabolic hubs where PA and other lipids can be selectively routed to mitochondrial membranes or LDs (Vance 2015, 2014;

Horvath and Daum 2013; Monteiro-Cardoso and Giordano 2024). However, lipid transport from mitochondria to MAM or LDs remains poorly understood. Here we show that ORP5 directs PA transport from mitochondria to MAM and also stabilizes it at MAM–LD junctions, promoting anabolic lipid storage in LD-associated mitochondria while limiting cardiolipin overproduction, thus coordinating storage-anabolic versus oxidative-catabolic mitochondrial programs.

Our lipidomics analysis further revealed that ORP5 deficiency alters specific TAG and cardiolipin species, rather than exerting a global effect on neutral or mitochondrial lipids (Fig. 5 A; Supp. Fig. 7). This specificity may reflect substrate selectivity of the enzymatic pathways that act downstream of PA partitioning. For example, mitochondria require PA with defined acyl chain compositions to generate mature cardiolipin species critical for respiratory chain assembly and cristae maintenance (Pennington et al. 2019). In parallel, defined TAG species contribute to LD dynamics and metabolic signaling (Wang 2015; Vanni et al. 2019). By tuning the relative flux of PA into these two pathways, ORP5 could thereby exert a broader influence on both mitochondrial function and systemic lipid homeostasis.

The redistribution of PA toward mitochondria upon ORP5 loss of function provides a mechanistic explanation for the mitochondrial hyperfusion phenotype we observed. Elevated mitochondrial PA is known to activate mitofusins and inhibit the fission factor DRP1, thereby shifting mitochondrial dynamics toward fusion (Kameoka et al. 2018). Concomitantly, we observed increased cardiolipin levels, especially of defined species such as cardiolipin 68:4 and 68:5, which contain combinations of C16 and C18 saturated and monounsaturated fatty acids (Fig. 5C). Cardiolipin has been shown to engage OPA1 to promote fusion and cristae maintenance. Additionally, cardiolipin with saturated acyl chains can block DRP1 oligomerization and therefore, prevent mitochondrial fission (Kameoka et al. 2018; Liu and Chan 2017). Thus, in ORP5-deficient cells, the combined accumulation of PA / cardiolipin and remodeled cardiolipin species may act synergistically to favor mitochondrial fusion and suppress mitochondrial division, leading to elongated mitochondria.

Notably, in ORP5-deficient cells, despite increased cardiolipin content, mitochondrial respiration was impaired (Monteiro-Cardoso et al. 2022), suggesting that excess cardiolipin with suboptimal acyl composition may compromise respiratory chain assembly rather than enhance it. Interestingly, deleterious effects of increased cardiolipin levels, including cardiolipin 68:5, have been associated to defective oxidative phosphorylation activity in MEGDEL syndrome, a rare neurometabolic disorder (Wortmann et al. 2012). These findings highlight the importance of ORP5 in safeguarding the balance between lipid-driven mitochondrial dynamics and respiratory capacity.

In 3T3-L1 white adipocytes, ORP5 is required for efficient PA channeling to LD and TAG accumulation (Fig. 1O-Q), suggesting that its role extends to specialized energy-storing tissues. ORP5 previously reported PS–PI(4)P exchange at ER–LD contacts (Du et al., 2019) may overlap with its newly described PA channeling function. An intriguing possibility is that PA transfer by ORP5 contributes to the regulation of PI(4)P levels at LDs, either independently or in parallel with PS exchange at MAM–LD contacts. Consistent with this, we observed a reduction in ORP5 protein levels at day 4 of adipocyte differentiation, which coincides with a decrease in PI(4)P recently reported (Wu et al. 2025). Wu and colleagues showed that LD PI4P recruits CIDE proteins to drive the transition from multilocular to unilocular LDs during adipogenesis and hepatic steatosis. Together with our findings, this underscores the importance of lipid signals at organelle interfaces in dictating LD identity and function. We propose a coordinated model: ORP5 ensures the upstream supply of PA for TAG synthesis at MAM–LD contacts, whereas PI(4)P–CIDE interactions subsequently organize LD size and morphology. Such multilayered regulation may be particularly relevant in adipocytes and hepatocytes, where imbalances in LD number or size directly impact systemic energy balance and disease states such as obesity and steatosis.

Several open questions emerge from our study. First, how ORP5 itself is regulated remains unknown. Future studies should address whether ORP5 is regulated by lipid availability, post-translational modifications, or dynamic protein–protein interactions at contact sites. ORP5 interacts with the mitochondrial tether PTPIP51, implicated in PA transport and monolysocardiolipin metabolism (Yeo et al. 2021) and it will be important to determine whether ORP5 activity is influenced by PTPIP51 abundance or local lipid flux. Second, although other lipid transfer proteins have been mainly studied at individual organelle contacts, such as VPS13 at ER–mitochondria or ER-LD contacts (Kumar et al. 2018), PTPIP51 at ER– mitochondria contacts (De Vos et al. 2012; Stoica et al. 2014; Yeo et al. 2021), and ORP2 at ER-LD contacts (Kentala et al. 2018), none have been thoroughly examined at the tripartite MAM-LD interface. MIGA2, which bridges mitochondria and LDs with the ER and promotes de novo lipogenesis in adipocytes (Freyre et al. 2019), represents one candidate, but whether and how it cooperates with ORP5 remains unknown. Our findings therefore position ORP5 as one of the few proteins with a defined role at these tripartite sites, though whether it acts independently or in concert with other LTPs remains to be established. Third, the contribution of ORP5 dysfunction to human metabolic disorders remains to be explored. Because disruptions in PA flux at MAM–LD contacts are predicted to simultaneously impair LD biogenesis and mitochondrial lipid metabolism, they could underlie pathological features of obesity, lipodystrophy, and fatty liver disease.

In conclusion, we identify ORP5 as a PA-specific transfer protein that safeguards mitochondrial lipid balance by preventing cardiolipin overload. At the MAM–LD interface, ORP5 redirects PA away from excessive cardiolipin synthesis toward TAG formation, thereby coupling lipid storage with mitochondrial integrity (Supp. Fig. 9G). Through this selective PA partitioning, ORP5 maintains lipid and energy homeostasis and coordinates metabolic fluxes across the mitochondria–ER–LD axis. These findings establish PA transport at tripartite contact sites as a central mechanism of inter-organelle lipid regulation and highlight the pivotal role of lipid transfer proteins in protecting mitochondrial function while tuning cellular energy metabolism.

## LIMITATION OF THE STUDY

Our study identifies ORP5 as a regulator of PA flux between mitochondria and LDs, balancing TAG storage and mitochondrial lipid metabolism, but some limitations still remain. First, while lipidomics revealed that ORP5 loss preferentially alters subsets of TAG and cardiolipin species, the molecular basis for this apparent acyl chain specificity remains unresolved. Whether this reflects intrinsic selectivity of the ORD domain for distinct PA species (as ORP8 displays acyl-selective activity for PS transfer (Ikhlef et al. 2021) or arises indirectly through remodeling enzymes that act downstream of ORP5-mediated transport, could not be addressed in this study. Also, the structural basis of PA recognition by ORP5 remains unresolved. High-resolution structural and biochemical studies will be essential to elucidate how ORP5 discriminates PA from other phospholipids, and how this activity is integrated into the dynamic architecture of organelle contacts. Second, although our *in vitro* assays establish that ORP5 can directly transfer PA, and our cell-based experiments show that it stabilizes PA at MAM–LD interfaces, most likely other MCS-resident proteins or metabolic enzymes contribute to shaping PA flux. Finally, while we observed that ORP5 deficiency alters mitochondrial morphology in parallel with increased cardiolipin, the precise causal links between altered lipid partitioning, mitochondrial dynamics, and organelle function were not directly dissected here. Future studies combining lipid flux tracing with high-resolution imaging of mitochondrial dynamics will be important to connect ORP5-mediated lipid transport to mitochondrial dynamics and functions.

## MATERIAL AND METHODS

### Antibodies, reagents and chemicals

Primary antibodies: rabbit anti-ORP5 (SIGMA, HPA038712), mouse anti-OPA1(BD Transduction Laboratories, 612607), mouse anti-DRP1 (BD Transduction Laboratories, 611113), rabbit anti-MFN1 (Proteintech, 13798-1-AP), rabbit anti-MFN2 (Proteintech, 12186-1-AP), rabbit anti-MFF (Proteintech, 17090-1-AP), mouse anti-GAPDH (GeneTex, GTX627408), mouse anti-Cytochrome C (BD Pharmingen, #556433), mouse anti-IP3R-3 (Transduction Laboratories, #610312), rabbit anti-Perilipin-3 (PLIN3, SIGMA, HPA006427), mouse anti-PDI (GeneTex, #GTX30716), mouse anti-GAPDH (GeneTex, #GTX627408) Secondary antibodies: anti-mouse-HRP (#NA931V) and anti-rabbit-HRP (# NA934V) from GE Healthcare. Lipid droplets markers: LipidTox deep Red (Invitrogen, H34477) and LipiBlue (Dojindo, LD01-10). Mitochondria markers: Mitotracker Orange CMTMRos (Invitrogen, M7510), Mitotracker Deep Red FM (Invitrogen, M22425) and Mitotracker Green FM (Invitrogen, M7514). Cell culture reagents: DMEM (Gibco, 31966), penicillin/streptomycin (Gibco, 15140-122), Opti-MEM (Gibco, 51985), non-essential amino acids (Gibco, 11140), Insulin (Merk, 19278), IBMX (#I5979, SIGMA), and dexamethasone (#D4902, SIGMA), collagen IV (#C5533, SIGMA).

Transfectant reagents: Lipofectamine 2000 (Invitrogen, 11668019), Lipofectamine 3000 (Invitrogen), Oligifectamine (Invitrogen, 11668019), Lipofectamine RNAi max (Invitrogen, 13778150).

Lipids with the following references were purchased from Avanti Polar Lipids: 1-oleoyl-2-(6-((4,4-difluoro-1,3-dimethyl-5-(4-methoxyphenyl)-4-bora-3a,4a-diaza-s-indacene-2-propionyl)amino)hexanoyl)-sn-glycero-3-phosphate (TopFluor-TMR-PA, #810240C), 23-(dipyrrometheneboron difluoride)-24-norcholesterol (TopFluor-Cholesterol, #810255P), 1-palmitoyl-2-oleoyl-sn-glycero-3-phosphocholine (POPC, #850457C), L-α-phosphatidylethanolamine, transphosphatidylated (Egg Trans PE, #841118C) 1,2-dioleoyl-sn-glycero-3-phosphoethanolamine-N-(cap biotinyl) (Biotinyl Cap PE, #870273C), 1-palmitoyl-2-(dipyrrometheneboron difluoride)undecanoyl-sn-glycero-3-phosphoethanolamine (TopFluor-PE, #810282C), 1-palmitoyl-2-(dipyrrometheneboron difluoride)undecanoyl-sn-glycero-3-phospho-L-serine (TopFluor-PS, #810283C). Lipin-1! inhibitor, propranolol hydrochloride (#P0884) was from Sigma-Aldrich.

### Plasmids

EGFP-ORP5A was described in (Galmes et al. 2016). EGFP-ORP5B, EGFP-ORP5DCC and EGFP-Opi1pQ2S-PABD were described in *(Guyard et al. 2022)*, and GFP-Sec22b in (Gallo et al. 2020). GST-ORD5 was described in (Monteiro-Cardoso et al. 2022). Mito-BFP (#49151) and ERoxBFP (#68126) were purchased from Addgene. Tom20-StayGold was a kind gift from Julien Prudent (Cambridge University, United Kingdom).

### Cell lines and culture conditions

HeLa *WT* and *ORP5KO* cells were acquired from Abcam (*WT*: ab255928; *ORP5KO*: ab265771), seipin KO cells were a kind gift from Hongyuan Yang (Yan et al. 2018). Adipocytes 3T3-L1 were provided by Xavier Prieur (Université de Nantes, France). HeLa cells were cultured at 37°C and 5% CO_2_ in DMEM containing 4.5 g/L glucose and supplemented with 10% FBS, 1% penicillin/streptomycin, and 1% non-essential amino acids. 3T3-L1 cells were cultured in Medium A, which consists of DMEM containing 4.5 g/L glucose and glutamax, and supplemented with 10% calf serum, 1% penicillin/streptomycin. For differentiation, cells were seeded at a density of 20,000 cells/mL in Medium B (same DMEM and supplement formulation of Medium A but with 10% fetal calf serum, instead of calf serum). Two days after confluence, adipocyte differentiation was induced in Medium B containing 2 µM insulin, 500 mM IBMX, and 1 µM dexamethasone. Three days after, the medium was changed to Medium B containing 2 µM insulin. On the sixth day of differentiation, cells were grown in Medium B. 3T3-L1 were used for experiments 10 days post differentiation.

### Transfections and siRNA

#### HeLa cells transfection

Cells were transfected with the indicated plasmids using Lipofectamine 2000 according to the manufacturer’s protocol. Briefly, HeLa cells were seeded in 13 mm diameter and 1.5 thickness circular coverglasses (Agar Scientific) and at 80% confluence cells were transfected with 1 mg of DNA for 3 h in serum depleted medium (Opti-MEM). The assays were performed 16 - 18 h post-transfection.

#### HeLa cells siRNA treatments

ORP5 and ORP8 knockdown in HeLa cells was achieved by siRNA transfection using Oligofectamine (Invitrogen) following the manufacturer’s instructions. Briefly, HeLa cells at 40% confluence, seeded in 13 mm 1.5 coverglasses (for imaging), 15 cm (for lipid droplets purification) or in 10 cm (for protein extract) tissue dishes, were transfected with 200 nM siRNA. Cells were cultured for 48 h prior additional treatment. Double-stranded siRNAs were derived from the following references: OSBPL5 Dharmacon, J-009274-10; OSBPL5 Dharmacon, J-009274-11; OSBPL8 Dharmacon, J-009508-06; OSBPL8 Dharmacon, J-009508-05 (Galmes et al. 2016).

#### 3T3-L1 Transfection

Adipocytes were transfected with the indicated plasmid for 4 hours in differentiation medium (DMEM + 10% FCS) without antibiotics using Lipofectamine 3000 according to the manufacturer’s protocol. Briefly, 1 hour before transfection, the cells were placed in differentiation medium without antibiotics. Just before transfection, glass coverslips placed in a 24-well plate were coated with type IV collagen (Sigma-Aldrich) at a concentration of 50 µg/mL for 30 min. Meanwhile, the mixtures containing the plasmid or Lipofectamine 3000 were prepared and incubated for 5 min at room temperature, while the mixture containing Lipofectamine was vortexed for a few seconds before incubation. The mixtures were then mixed and incubated for 15 min at room temperature. During incubation, the adipocytes were detached from the Petri dish using Tryple (LifeTechnologies) and resuspended in a differentiation medium without antibiotics. The adipocytes were then seeded on glass slides after adding the mix containing the transfected plasmid. The cells were imaged approximately 18 hours after transfection.

#### 3T3-L1 siRNA treatment

At day 6 of differentiation, cells were dissociated with TrypleE express (Thermofisher) and resuspended in medium B. The siRNA mix (20 nM siRNA, 2.5 μL Lipofectamine RNAi max, Thermofisher, 47.5 μL of OptiMEM, Thermofisher), was added to a collagen IV precoated 13 mm coverslips or 10 cm tissue culture dishes, and 0.25 mL of differentiated adipocyte suspension is added for reverse transfection. 72 hours after, the assays are performed. Double-stranded siRNA targeting mouse ORP5 was obtained from Dharmacon, reference J-046295-11.

##### RT-qPCR

Total RNA was extracted from 3T3-L1 cells using the RNeasy mini kit (Qiagen, Valencia, CA). Complementary DNA was generated from 1 μg of RNA using random hexamers and the QuantiTect Reverse Transcription Kit (Quiagen, #205311) following manufacturer’s instructions. Quantitative real-time PCR was performed using LightCycler^®^ 480 SYBR Green I Master (Roche, #04707516001) in an LightCycler^®^ 96 Instrument Real-Time System (Roche). Reactions with 1 ng complementary DNA were performed in three technical replicates using the following mouse primers: 18S (Teixeira et al., 2022) Fw: ACCGCAGCTAGGAATAATGGA; Rv: GCCTCAGTTCCGAAAACCA; PPARγ (Loving et al., 2021) Fw: GACGCGGAAGAAGAGACCTG; Rv: GGAATGCGAGTGGTCTTCCA. Gene expression analyses were performed by relative quantification using *18S* as the house-keeping control, using the comparative ΔΔC_t_ method (Schmittgen and Livak 2008).

### Immunofluorescence

3T3-L1 were seeded on 13 mm glass bottom coverslips (Agar Scientific). At 10 days of diffrention 3T3-L1 adipocytes were incubated with 1 μM MitoTracker (mitochondrial marker, Invitrogen) and Lipiblue for 30 min at 37°C, 5% CO_2_ and then fixed with 4% PFA/PBS for 30 min at room temperature. Fixed cells were then washed in PBS and incubated with 50 mM NH4Cl/PBS for 15 min at room temperature. After washing with PBS and blocking buffer (2% BSA, 0.1% Saponin in 1xPBS), cells were incubated with primary antibodies rabbit anti-ORP5 (1:150) and rabbit anti-ORP8 (1:150) diluted in blocking buffer for 1 h at room temperature and then with fluorescently-labeled secondary Alexa Fluor 488 anti-rabbit (1:500, Invitrogen, A-11008). After washing with blocking buffer and then PBS, coverslips were mounted on microscopy slides and images were acquired on Confocal inverted microscope SP8-X (DMI 6000 Leica). Images from a mid-focal plane are shown. Images were processed and fluorescence was analyzed off line using Image J.

### LD Biogenesis

LD biogenesis HeLa cells were plated in DMEM containing 5% of lipoprotein depleted serum (SIGMA) to induce the consumption of pre-existing lipid droplets. LD biogenesis rescue experiments were conducted in control and knockdown for ORP5 cells expressing Mito-BFP together with EGFP-ORP5B, EGFP-ORP5B L489D or EGFP-Sec22b. Lipid droplet formation was induced by treating cells with 1 μM FA^568^ for 15 min at 37°C and 5% CO_2_. Cells were fixed in 4% PFA and lipid droplets were stained with LipidTox (1:1000) in 1xPBS for 30 min. Newly formed LDs marked simultaneously by LipidTox and FA^568^ were counted using Image J software.

### Chemical inhibition of Lipin

Lipin-1! inhibition was achieved by treating cells with propranolol hydrochloride. Briefly, cells were incubated with 100 µM propranolol hydrochloride in DMEM containing 4.5 g/L glucose supplemented with 10% FBS and 1x nonessential amino acids for 1 h at 37°C and 5% CO_2_.

### Incorporation of TopFluor lipids in HeLa and 3T3L-1

TopFluor lipids were incorporated in cells as previously described (Sassano et al. 2023) with slight modifications. Briefly, TopFluor-TMRA-PA and TopFluor-Cholesterol in chloroform were dried using argon gas. TopFluor-TMRA-PA and TopFluor-Cholesterol were then resuspended in ethanol to a final stock concentration of 0.5 M. For cell treatment 2 uM TopFluor lipids were complexed with 2% FBS in 1xPBS. HeLa and 3T3L-1 were incubated with TopFluor / FBS complex for 30 min at 37°C and 5% CO_2_.

### Cell swelling

After transfection the cultured cells were transferred into a hypotonic culture media DMEM:H2O (5:95% v/v) at pH 7.4, at 37 °C, 5% CO₂, for 15 min, to induce cell volume increase and Giant Organelle Vesicules (GOVs).

### Microscopy

#### Confocal

HeLa and 3T3-L1 were stained with 1 μM Mitotracker and/or 0.1 μM LipiBlue in serum free media Opti-MEM for 30 min at 37°C and 5% CO_2_. Cells were then fixed with 4% PFA in 1xPBS for 30 min at room temperature, washed in 1xPBS and then incubated with 50 mM NH_4_Cl in 1x PBS for 15 min at room temperature. After washing with 1xPBS, cells (which were not stained with Lipiblue) were incubated with LipidTox deep red (1:1000) in 1xPBS to stain lipid droplets. Finally, coverslips were mounted on microscopy slides using Vectashield (Vector Laboratories, H-1000-10) and sealed with nail polish. Images were acquired on an inverted confocal microscope SP8-X (DMI 6000 Leica). Optical sections were acquired with a Plan Apo 63x oil immersion objective (N.A. 1.4, Leica) using the LAS-X software. Fluorescence was excited using either a 405nm laser diode or a white light laser, and later collected after adjusting the spectral windows with GaAsP PMTs or Hybrid detectors. Images from a mid-focal plane are shown. Images were processed and fluorescence was analysed offline using Image J and IMARIS. The number and volume of LDs as well as PA-derived signal in LDs was assessed using Imaris. LDs were detected and modulated using the surfaces option in IMARIS. Then using Imaris statistics tool we obtained the number of LDs per cell, their volume and PA mean intensity. To avoid differences caused by distinct uptake of TopFluor-TMR-PA, PA mean intensity in LDs was normalized by the PA mean intensity in the corresponding cells. The results were expressed in function of the control. The LD/mitochondria TopFluor-TMR-PA intensity ratio was determined by plotting fluorescence profiles in regions where LDs were closely associated with mitochondria. The mean TopFluor-TMR-PA intensity within LDs was divided by the corresponding mean intensity within mitochondria. For mitochondria morphology analysis, aspect ratio (AR) and form factor (FF) which are indicators of the mitochondrial length and branching, respectively, have been quantified using ImageJ software as in (Chen et al. 2021). Briefly, the images were convolved using the matrix developed by Koopman (Koopman et al. 2016; Marchi et al. 2017), then using the threshold tool mitochondria were isolated from the background.

#### Live imaging

HeLa *WT* and *ORP5KO* cells were seeded on glass bottom Ibidi chambers (µ-slide 2 wells) 2 days prior imaging. Cell imaging was performed on an inverted Nikon Ti Eclipse E microscope coupled with a Spinning Disk (CSU-X1-A1; Yokogawa) and cage incubator to control both temperature and CO_2_ (37°C, 5% CO_2_). After excitation at 488 nm 561 nm and 642 nm, fluorescent signals were detected with a 40× oil immersion objective (PLAN FLUOR; NA: 1.30; Nikon), an emission filter Quad bandpass 525/45, 590/20, 692/40 nm (Semrock), and a Prime 95B sCMOS camera (Photometrics). Images were captured every 20 s for 30 min. Approximately 1 min after the start of captures, TopFluor-TMR-PA / FBS complex was added to a final concentration of 2 µM. For *ex vivo* and *in situ* cell swelling live-cell imaging, the images were acquired using a Carl Zeiss LSM800 microscope with an oil-immersed ×63 objective. Images of LDs (from Bodipy) and ER vesicles were segmented using custom-trained StarDist models on Python (v3.10)(M. Weigert 2020). Image analysis was performed on CellProfiler (v4.2.6). PA signal was measured using the “MeasureObjectIntensity” module in LDs mask and from the edges of ER vesicles (2-pixel inside + 2-pixel outside the border of the masks). Distribution of PA signal (%PA in LDs) between organelles was calculated by dividing the PA signal in LDs mask by the sum of PA signal from both LDs and ER.

#### Electron Microscopy

HeLa cells expressing HRP-KDEL were fixed on coverslips with 1.3% glutaraldehyde in 0.1 M cacodylate buffer, washed in 0.1 M ammonium phosphate [pH 7.4] buffer for 1 h and HRP was visualized with 0.5 mg/mL DAB and 0.005% H_2_O_2_ in 0.1 M Ammonium Phosphate [pH 7.4] buffer. Development of HRP (DAB dark reaction product) took between 5 min to 20 min and was stopped by extensive washes with cold water. Cells were postfixed in 2% OsO_4_+1% K_3_Fe(CN)_6_ in 0.1 M cacodylate buffer for 1 h at 4°C, washed in cold water and then contrasted in 0.5% uranyl acetate for 2 h at 4°C, dehydrated in an ethanol series and embedded in epon as for conventional EM. Ultrathin sections were counterstained with 2% uranyl acetate and observed under a FEI Tecnai 12 microscope equipped with a CCD (SiS 1kx1k keenView) camera.

### Cell fractionation

#### Lipid droplets

For lipid droplets purification, lipid droplets were induced in HeLa cells (80x10^6^ cells) with 250 uM oleic acid treatment overnight. HeLa cells were harvested 48 hours after transfection with siRNA oligos and washed with 1xPBS by centrifugation at 600 *g* for 5 min. Cell pellet was then resuspended in buffer A (225 mM mannitol, 75 mM sucrose and 30 mM Tris-HCl, 0.5 mM EGTA, pH 7.4) and homogenized using a Tissue Grinder Dura-Grind®, Stainless Steel, Dounce (Wheaton). The homogenate was centrifuged at 1000 *g* for 10 min to remove nuclei and unbroken cells. Transfer the supernatant to an ultracentrifuge tube and fill it up with buffer B (20 mM HEPES, 100 mM KCl and 2 mM MgCl2, pH 7.4). Centrifuge at 200 000 *g* for 1h30 at 4°C. Carefully collect lipid droplets from the top band of the gradient formed and transfer them into an Eppendorf tube using a 200-μl pipette.

#### MAM and mitochondria

HeLa cells (150x10^6^ cells) were harvested 48 hours after transfection with siRNA oligos and washed with PBS by centrifugation at 600 g for 5 min. The cell pellet was resuspended in buffer A (225 mM mannitol, 75 mM sucrose and 30 mM Tris-HCl, 0.5 mM EGTA, pH 7.4) and homogenized using a glass teflon potter, as in (Galmes et al. 2016; Monteiro-Cardoso et al. 2022). The homogenate was centrifuged at 600 *g* for 5 min to remove nuclei and unbroken cells. The crude mitochondria were pelleted by centrifugation at 10 000 *g* for 10 min. To separate MAM and pure mitochondria fractions, the pellet was resuspended in MRB buffer (250 mM mannitol, 5 mM HEPES and 0.5 mM EGTA, pH 7.4) and layered on top of different concentrations of Percoll gradient (225 mM mannitol, 25 mM HEPES, 1 mM EGTA pH 7.4 and 30% or 15% Percoll). After centrifugation at 95 000 *g* for 30 min, two dense bands containing either the pure mitochondria or MAM fraction were recovered and washed with MRB buffer by centrifugation at 6300 *g* for 10 min to remove residual Percoll and residual contamination. MAM was pelleted by centrifugation at 100 000 *g* for 1 hour.

### Western blotting

For immunoblotting, total cells, MAM, mitochondria or LD were resuspended in ice cold lysis buffer [50 mM Tris, 150 mM NaCl, 1% Triton X-100, 10 mM EDTA, pH 7.2, and protease inhibitor cocktail (Roche). Lysates were then centrifuged at 21 000 g for 30 min at 4°C and protein concentrations were determined by Bradford assay. The supernatants were boiled in reducing the SDS sample buffer and 30 µg of protein were solved on 4-20% gradient SDS-PAGE gels (Biorad). The separated proteins were transferred to 0.45 μm Nitrocellulose membrane (GE Healthcare). The membrane was blocked by 5% non-fat milk in the TBS-T buffer (1xTBS buffer with 0.1% Tween 20) for 1 h at room temperature. Then the membrane was incubated with the primary antibodies at 4°C overnight, using the following dilutions: rabbit anti-ORP5 (SIGMA, 1:1000), rabbit anti-ORP8 (Gentex, 1:1000), mouse anti-Cytochrome C (BD Pharmingen, 1:2000), mouse anti-PDI (GeneTex, 1:500), rabbit anti-PLN3 (SIGMA, 1:800), mouse anti-IP3R-3 (BD Transduction Laboratories, 1:2000), mouse anti-DRP1 (BD Transduction Laboratories, 1:1000), rabbit anti-MFF (Proteintech1:1000), rabbit anti-MFN1 (Proteintech, 1:1000), rabbit anti-MFN2 (Proteintech, 1:1000), mouse anti-OPA-1 (BD Transduction Laboratories, 1:1000), mouse anti-GAPDH (GeneTex, 1:10000) in 5%milk in TBST. The membrane was then incubated with the Horseradish peroxidase-conjugated anti-rabbit IgG secondary antibody (1:5000, in 5%milk in TBST) or anti-mouse IgG secondary (1:5000, in 5%milk in TBST) at room temperature for 1h, followed with washing and detection using the enhanced chemiluminescence (ECL) detection kit (Cytiva). For Western blot quantification, bands of protein of interest were detected using ChemiDoc™ Imaging Systems (Life Science Research, Bio-Rad) and analyzed using Imaje J.

### Lipidomic analysis

*LC-MS-based lipidomics analysis of whole cells WT and ORP5KO and of lipid droplets isolated from control and RNAi treated HeLa Cells:*

Dried down samples were reconstituted in 80 μL methanol and incubated at 30°C for 5 mins with vigorous vortex. Samples (1 μL) were analyzed by LCMS (Agilent 1290 infinity/Infinity II Agilent) MS (Agilent 6495c triple quadruople). Acquisition DMRM (dynamic multiple reaction monitoring) method used was referred as described previously (Huynh et al. 2019) with a modification for some lipid species (Dass et al. 2024). The resulted LCMS data was subjected to targeted analysis using Mass Hunter Quantification software (Agilent). Each lipid species was quantified using a calibration curve of each representative lipid with known abundance. Then each lipid abundance was normalized according to the cell ratio.

*LC-MS-based lipidomics analysis of whole cells, MAM and mitochondria isolated from control and RNAi treated HeLa Cells:* Lipid extracts obtained from whole cells, MAM and mitochondria isolated from control and RNAi-treated cells were analyzed by Lipotype GmbH (Dresden, Germany) as a fee-for-service, as described below.

*Lipid extraction:* Mass spectrometry-based lipid analysis was performed by Lipotype GmbH (Dresden, Germany) as described. Lipids were extracted using a two-step chloroform/methanol procedure. Samples were spiked with internal lipid standard mixture containing: cardiolipin 16:1/15:0/15:0/15:0 (CL), ceramide 18:1;2/17:0 (Cer), diacylglycerol 17:0/17:0 (DAG), hexosylceramide 18:1;2/12:0 (HexCer), lyso-phosphatidate 17:0 (LPA), lyso-phosphatidylcholine 12:0 (LPC), lyso-phosphatidylethanolamine 17:1 (LPE), lyso-phosphatidylglycerol 17:1 (LPG), lyso-phosphatidylinositol 17:1 (LPI), lyso-phosphatidylserine 17:1 (LPS), phosphatidate 17:0/17:0 (PA), phosphatidylcholine 17:0/17:0 (PC), phosphatidylethanolamine 17:0/17:0 (PE), phosphatidylglycerol 17:0/17:0 (PG), phosphatidylinositol 16:0/16:0 (PI), phosphatidylserine 17:0/17:0 (PS), cholesterol ester 20:0 (CE), sphingomyelin 18:1;2/12:0;0 (SM), triacylglycerol 17:0/17:0/17:0 (TAG) and cholesterol D6. After extraction, the organic phase was transferred to an infusion plate and dried in a speed vacuum concentrator. 1st step dry extract was re-suspended in 7.5 mM ammonium acetate in chloroform/methanol/propanol (1:2:4, V:V:V) and 2nd step dry extract in 33% ethanol solution of methylamine in chloroform/methanol (0.003:5:1; V:V:V). All liquid handling steps were performed using Hamilton Robotics STARlet robotic platform with the Anti Droplet Control feature for organic solvents pipetting.

*MS data acquisition:* Samples were analyzed by direct infusion on a QExactive mass spectrometer (Thermo Scientific) equipped with a TriVersa NanoMate ion source (Advion Biosciences). Samples were analyzed in both positive and negative ion modes with a resolution of Rm/z=200=280000 for MS and Rm/z=200=17500 for MS/MS experiments, in a single acquisition. MSMS was triggered by an inclusion list encompassing corresponding MS mass ranges scanned in 1 Da increments (Surma et al. 2015). Both MS and MSMS data were combined to monitor CE, DAG and TAG ions as ammonium adducts; PC, PC O-, as acetate adducts; and CL, PA, PE, PE O-, PG, PI and PS as deprotonated anions. MS only was used to monitor LPA, LPE, LPE O-, LPI and LPS as deprotonated anions; Cer, HexCer, SM, LPC and LPC O-as acetate adducts and cholesterol as ammonium adduct of an acetylated derivative.

*Data analysis and post-processing:* Data were analyzed with in-house developed lipid identification software based on LipidXplorer (Herzog et al. 2011; Herzog et al. 2012). Data post-processing and normalization were performed using an in-house developed data management system. Only lipid identifications with a signal-to-noise ratio >5, and a signal intensity 5-fold higher than in corresponding blank samples were considered for further data analysis. The results were expressed as Mol% of total lipids detected in each sample.

### Lipid transfer Assay

*Purification of ORP5 ORD domain:* Purification of ORP5 ORD was achieved as previously described (Ref Cell Reports). Escherichia coli BL21DE3 RILP (Invitrogen) cells were transformed with plasmids encoding for GST tagged ORP5 ORD domain following the manufacturer’s instruction. Bacteria were then grown overnight at 37°C and used to inoculate a large-scale volume (1L). When the OD_600_ reached 0.4, cultures were cooled down and incubated at 18°C until they reached O_D600_ = 0.65. Cells were induced by addition of isopropyl β-D-1-thiogalactopyranoside to a final concentration of 0.1 mM and incubated overnight at 18°C before harvesting. Cells were resuspended in 35 mL binding buffer (1X PBS, 1 mM EDTA, 1 mM DTT, Protease inhibitor) then 250 units of benzonase nuclease (Sigma) were added to the resuspension. Cells were lysed by sonication and the supernatant was recovered after 20 min centrifugation at 184 000g and 4°C. Supernatant containing GST tagged proteins was incubated with 2 mL of Glutathione Sepharose 4 fast flow for 1 hour at 4°C under nutation. Beads were washed using a series of wash buffers: 1^st^ (1X PBS, 1 mM EDTA, 1 mM DTT), 2^nd^ HSP-removal buffer (50 mM Tris pH 7.5, 50 mM KCl, 20 mM MgCl_2_, 5 mM ATP) then cleavage buffer (50 mM Tris pH 7.5, 150 mM NaCl, 1 mM EDTA, 1 mM DTT). Cleavage of the GST tag was realized overnight at 4°C using Prescission protease. Untagged proteins were eluted with cleavage buffer, flash frozen and stored at -80°C until lipid transfer assay was performed.

*Liposome preparation:* 1 µmol of the appropriate lipid mixtures in chloroform solution was dried in a glass tube for 10 min under a gentle stream of argon, and then for 1 hour under vacuum. The dried lipid films were resuspended in 1 mL of buffer H (25 mM HEPES/KOH, pH 7.7; 150 mM KCl; 10%(v/v) Glycerol) by vigorously vortexing for 30 min at room temperature. Unilamellar liposomes were produced by seven freeze-thaw cycles (30 sec in liquid nitrogen followed by 5 min in a 37°C water bath) and extrusion (at least 21 times) through a polycarbonate filter with 100 nm pore size (polycarbonate membranes from Avanti Polar Lipids). The liposomes were then stored on ice.

*Lipid Transfer:* The lipid transfer assays were realized with liposomes as previously described. Briefly, the donor liposomes contained 1% mol TopFluor lipids (-PS, -PE or -PA) and 2% mol Biotinyl Cap PE. The acceptor liposomes contained 75% POPC and 25% Egg Trans PE or POPC only. For each reaction, 25 µL of streptavidin-coated magnetic beads (DynabeadsMyOne Streptavidin T1, Invitrogen) were washed in buffer H and mixed with 25 µL of 1 mM donor liposomes. The mixture was incubated for 1 hour at 25°C with intermittent gentle mixing. Bead-bound donor liposomes were then washed, resuspended in 25 µL and mixed with 25 µL of 1 mM acceptor liposomes and 50 µL of buffer H or protein (0.3 µM protein and 2.5 µM TopFluor lipids in the reaction solution). The mixture was incubated at 37°C for 1 hour with intermittent gentle mixing. Supernatant containing acceptor liposomes was recovered after binding of bead-bound donor liposomes to a magnetic rack. TopFluor fluorescence of acceptor and donor liposomes was measured (after solubilization with 0.4% (w/v) n-dodecyl-β-D-maltoside, DDM) in a SpectraMax M5 plate reader (Molecular Device) equilibrated to 30°C (TopFluor-PE/PS, excitation: 450 nm; emission: 510 nm; cutoff: 475 nm; low gain; TopFluor-TMR-PA, excitation: 510 nm; emission: 571 nm; cutoff: automatic; low gain). The percentage of lipids transferred from donor to acceptor liposomes was determined using the following formula: 100*F_acceptor_/(F_acceptor_+F_donor_).

### *Ex vivo* lipid transfer

HeLa WT cells were transfected with either an ER or mitochondria plasmid marker (ER: EGFP-ORP5; Mitochondria: Mito-BFP). The micromanipulation measurements were arranged as previously described by Santinho et al. (Nature Communications 2024(https://doi.org/10.1038/s41467-024-48086-7)). Briefly, cells were swollen with diluted aqueous DMEM (5:95% v/v; Dutscher) at 37°C, 5% CO2 for 15 min to create a hypotonic shock. Giant organelle vesicles were then extracted by mechanically lysing the cells. Donor GOVs were incubated for 30 min with TopFluor-TMR-PA (0,33 nM) dissolved in ethanol. Afterwards, the incubation media was exchanged to eliminate unbound PA. Donor and acceptor GOVs were placed on a BSA (10% v/v) pre-treated coverslip (Menzel Glaser; 24 × 36 mm, no. 1) in a 1:1 volume ratio.

The glass pipettes (pulled from borosilicate capillaries: 1.0OD, 0.58ID, 30-0017 GC100-15b; Harvard Apparatus, Holliston, MA) were cleaned utilizing a plasma cleaner, covered with mPEG5K-silane (Cat#JKA3037 Merck) and filled with diluted DMEM. The pipettes were placed in the homemade setup for the micromanipulation measurements and aligned with the confocal microscope settings. Donor and acceptor GOV were carefully captured with the micropipettes and placed apart without applying high aspiration on the GOV. The selected GOV pair was then brought in contact by moving the pipettes closer together. PA intensity was monitored via CLSM in 5-minute increments. For the intensity analysis of micrographs of the time increments and the contact area residue the software Fiji was used. The line profile tool was utilized to measure the PA intensity on GOV membranes at an equatorial location where neither the pipette nor the contact area was intervening.

Intensity values were background corrected and fitted to a Gaussian curve.

Calculation of the share of transferred PA from donor to acceptor organelle after 30 min: assuming no mass conservation, first the ratio of the sums of PA intensity in donor (*Int_donor_*) and acceptor (*Int_acc_*) GOVs at time point t=0 min and t=30 min was calculated, resulting in the normalization factor f.

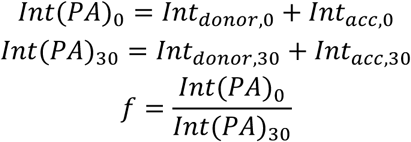

f was then applied to the share of transferred PA from donor to acceptor organelle after 30 min calculation as followed:

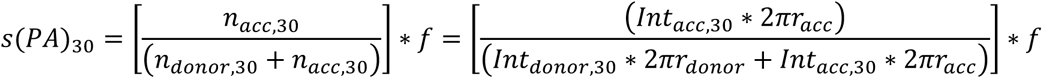

With *n*_30_ being the amount of PA in time frame 30 min. n for PA was obtained by utilizing the circumference formula of the spherical GOV, where 4 is the radius of the GOV, multiplied by the measured intensity *int*_30_. Abbreviation *acc* and *donor* indicate the values for acceptor and donor GOV, respectively.

### Statistical Analysis

The data were presented as the mean ± standard error of the mean (SEM) of at least three independent experiments. Boxplot representations indicate the first (Q1/25th percentile), median (Q2/50th percentile), and third (Q3/75th percentile) quartiles, and whiskers are a maximum of 1.5 times the interquartile range obtained at least from three independent experiments. Data points beyond the 5th and 95th percentiles were plotted individually as outliers, and the mean was marked with a cross (+). Data derived from different groups were compared by Student’s *t* test, Kruskal-Wallis test, or Mann-Whitney U test. All statistical analyses were performed using GraphPad Prism software. *p < 0.05, **p < 0.01, ***p < 0.001, and ****p < 0.0001 were considered significant.

## ACKNOWLEDGEMENTS

The authors would like to thank the group members for helpful discussions, Juan Reguera for insightful input, and Xavier Prieur for valuable discussions and for sharing adipocyte-handling protocols. We would like to thank Gaelle Dufayet-Chaffaud for technical assistance and Angelique Nicolas for help in ordering and administrative tasks essential for this work. We apologize to all whose work was not cited due to space constraints. This work was supported by the Agence Nationale de la Recherche (ANR-22-CE11-0024-01/MADE to FG/ART and ApicoLIpidTraffic to CB/FG/YYB), the the Association Française contre les Myopathies (AFM-Téléthon Research Grant 23778 to DT/FG), add CNRS-MITI to FG/ART and FA to FG. The present work has benefited from Imagerie-Gif core facility supported by l’Agence Nationale de la Recherche (FBI ANR-24-INBS-0005 (BIOGEN); SPS ANR-17-EUR-0007, EUR SPS-GSR). CB, YYB, and the GEMELI lipidomics platform were supported by Agence Nationale de la Recherche, France (Project ApicoLipidAdapt grant ANR-21-CE44-0010 ; Project Apicolipidtraffic grant ANR-23-CE15-0009-01; Project OIL grant ANR-24-CE15-2171-02; Project PlasmoHost ANR-25-CE30), The Fondation pour la Recherche Médicale (FRM EQU202103012700), Laboratoire d’Excellence Parafrap, France (grant ANR-11-LABX-0024), LIA-IRP CNRS Program (Apicolipid project), the Université Grenoble Alpes (IDEX ISP Apicolipid), Région Auvergne Rhone-Alpes (Grant IRICE Project GEMELI), Infrastructures en Biologie Santé et Agronomie (IBiSA), and the Collaborative Research Program Grant CEFIPRA (MESRI-DBT, Project 6003-1, and Project 7152). We also acknowledge the ImagoSeine core facility of Institut Jacques Monod, member of France-BioImaging (ANR-10-INBS-04) and IBiSA, with the support of Labex “Who Am I”, Inserm Plan Cancer, Region Ile-de-France and Fondation Bettencourt Schueller.

## AUTHORS CONTRIBUTION

VFMC, ART and FG designed the research. VFMC, VG, JT, CS, HE, YYB, CB, MZ, ART and FG designed experiments. VFMC conducted all the *in situ* experiments with assistance from VG, who carried out the experiments on seipin KO cells and adipocytes. CS performed the mitochondria and MAM purifications. CB and YYB conducted and analyzed the lipidomic experiments on total cells lipid extracts. DT assisted for the *in vitro* lipid transfer experiments. NEK performed the electron microscopy analysis and assisted VFMC in the LD purification experiments. HE, MZ performed some of the *in situ* swelling and GOV experiments. JT performed the *ex vivo* GOV experiments. VFMC and FG wrote the manuscript, which was reviewed and edited by all co-authors.

---

**Supplementary Figure 1:**
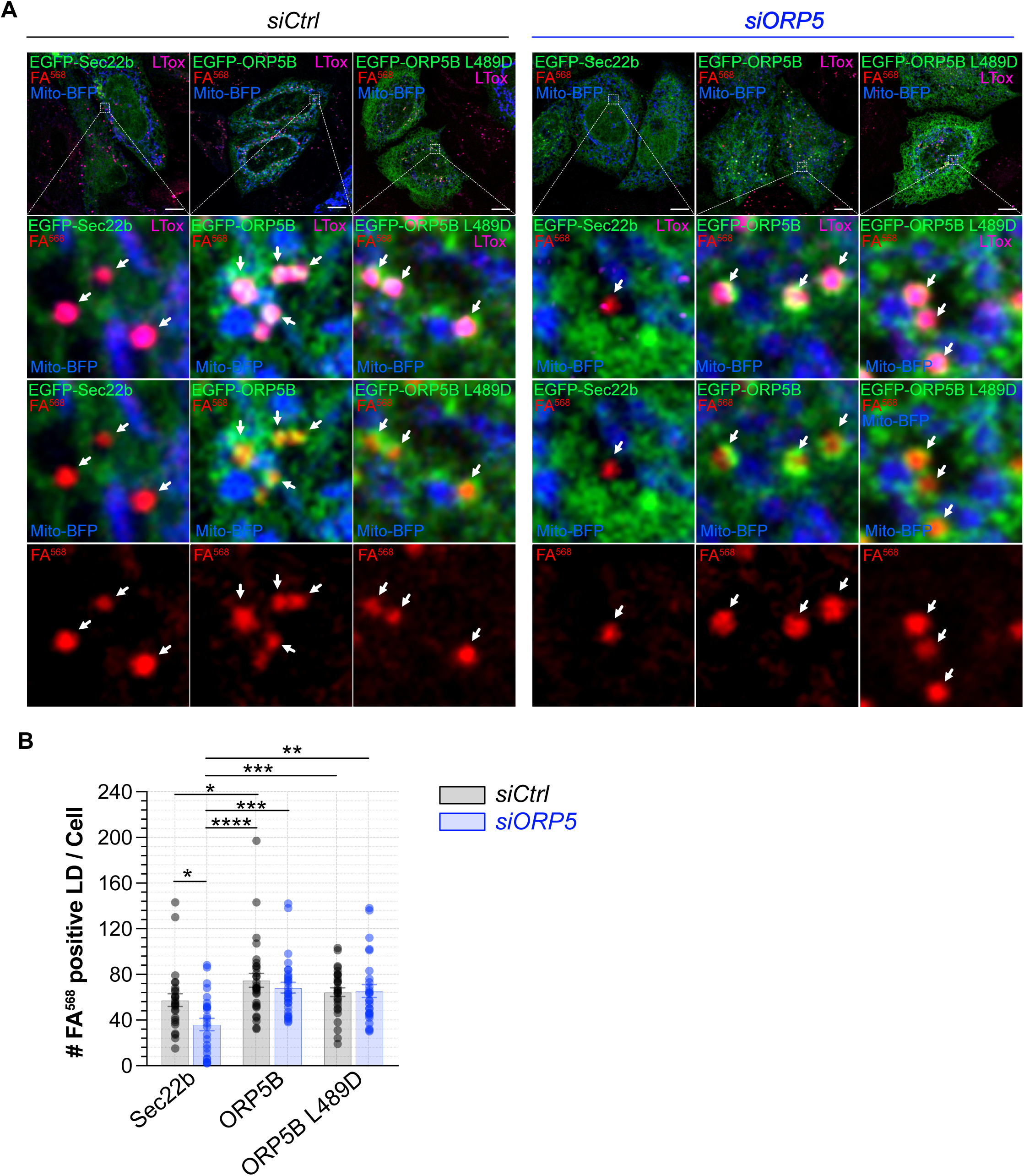
**(A)** Confocal (single focal plane) micrographs of control and ORP5 knockdown delipidated HeLa cells co-overexpressing Mito-BFP (blue) with EGFP-Sec22b (green), EGFP-ORP5B (green), EGFP-ORP5B L489D (green). Arrowheads indicate the newly formed LD. Scale bar, 10 μm. **(B)** Quantitative analysis of the number of FA^568^-positive LD in control and ORP5 knockdown delipidated HeLa cells co-overexpressing Mito-BFP and Sec61β-EGFP, siRNA-resistant EGFP-ORP5B (green), EGFP-ORP5B L489D, and treated for 15 with FA^568^. Data are shown as mean the number of newly formed lipid droplets ± SEM of *n* = 26 - 31 cells. Dara were compared using Kruskal–Wallis test followed by multiple comparisons, *p < 0.05,**p < 0.01, \*\*\**p* < 0.001, ****p < 0.0001.

**Supplementary Figure 2:**
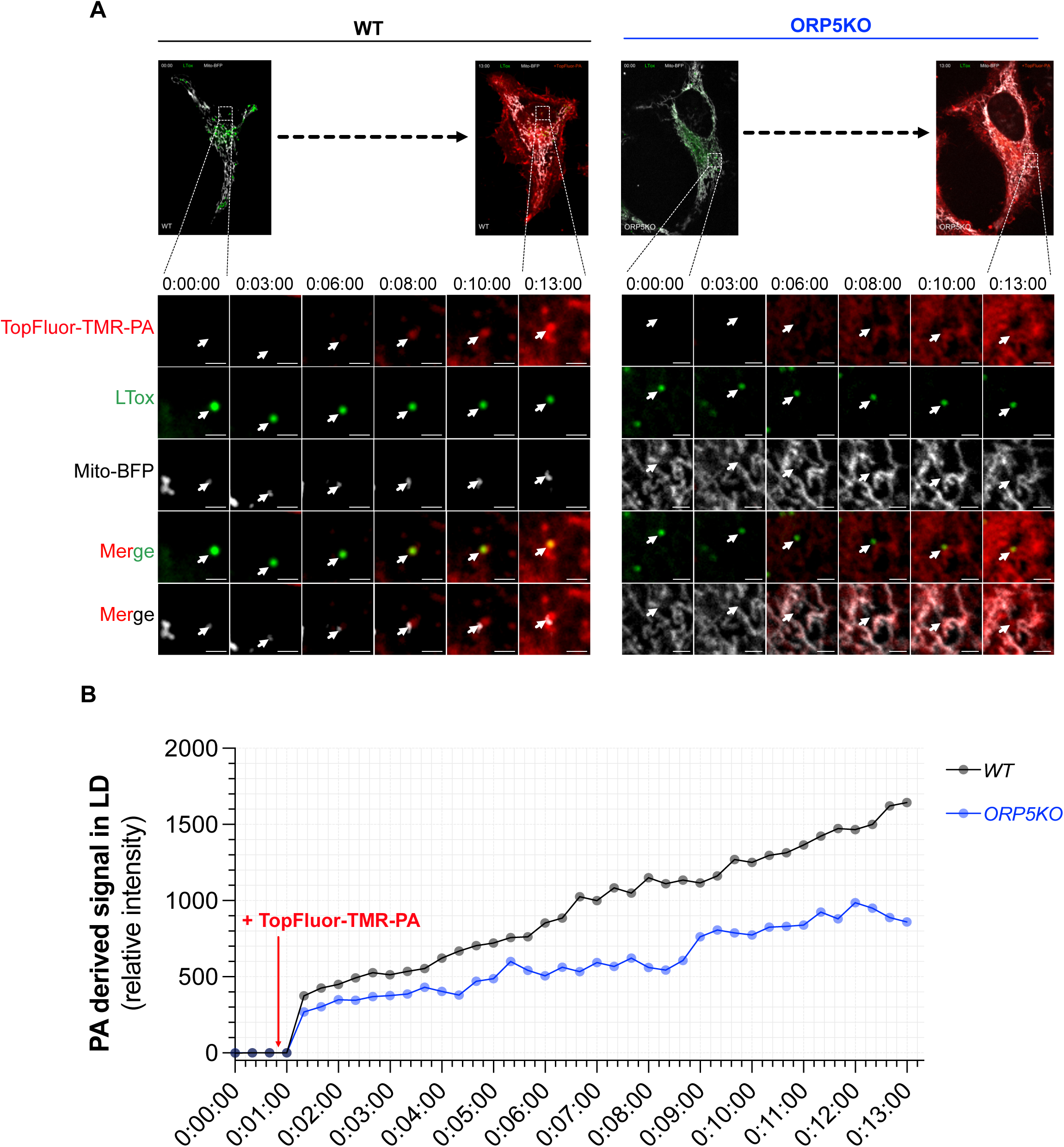
**(A)** Spinning video snapshots of HeLa *WT* and *ORP5KO* cells stained with mitotracker green (gray) and LipidTox (green) and treated with 2 µM TopFluor-TMR-PA (red). After 40 seconds of acquisition, the cells were treated with TopFluor-TMR-PA. Arrows indicate regions where TopFluor-TMR-PA accumulates in co-localization with lipid droplets. Scale bar, 1 µm. **(B)** Intensity profile changes of TopFluor-TMR-PA associated with lipid droplets during 13 minute recordings.

**Supplementary Figure 3:**
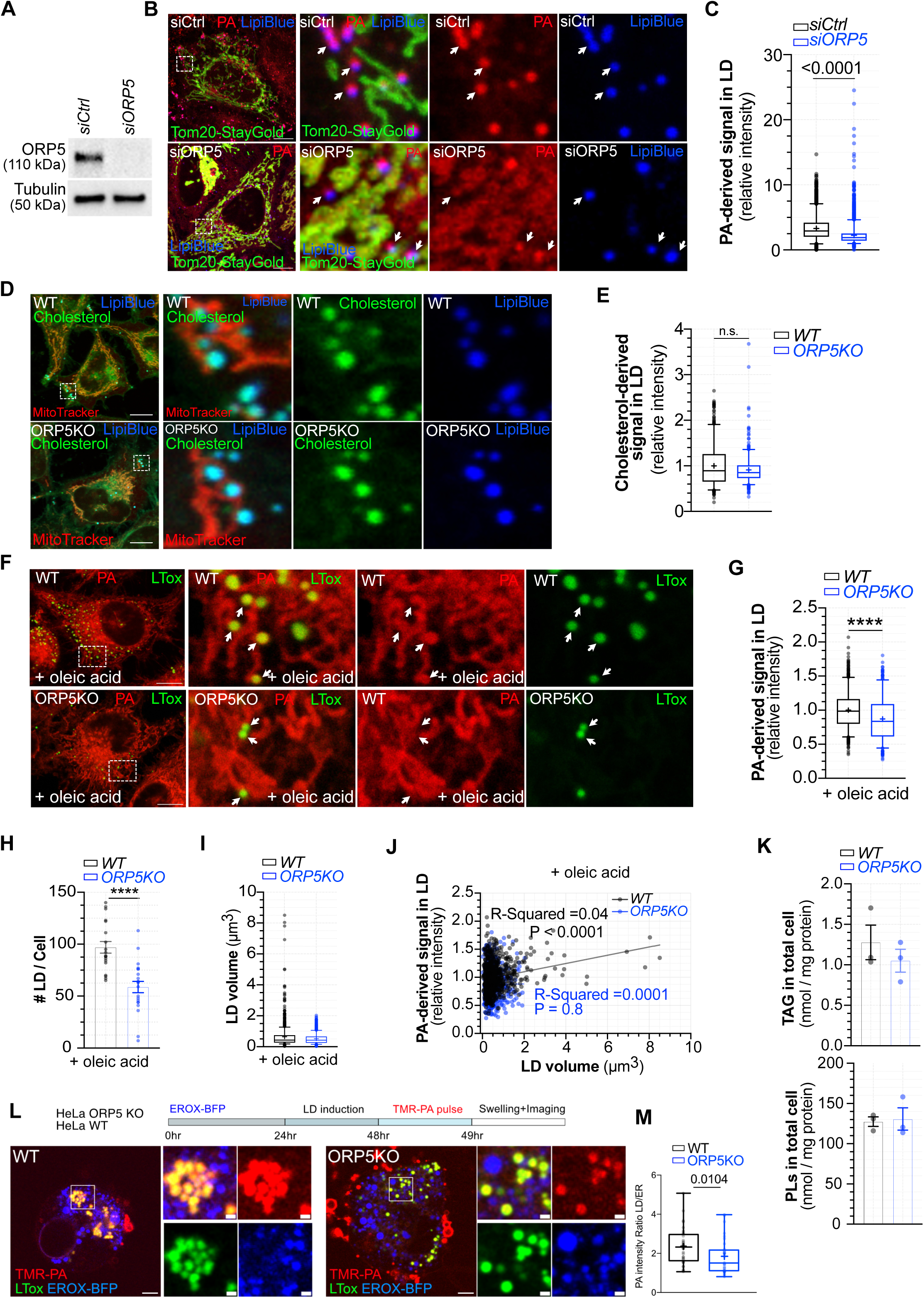
**(A)** Representative immunoblotting of RNAi-targeting ORP5 efficiency in reducing ORP5 expression. **(B)** Representative confocal images, including full cell and zoomed regions, of HeLa cells expressing Tom20-StayGold (mitochondria, green) and treated with 2 µM TopFluor-TMR-PA (red) for 30min 48 h after transfection with control siRNA (*siCtrl*) or with a *ORP5*-specific siRNA (*siORP5*). Lipid droplets were stained with Lipiblue (blue). Scale bar, 10 µm. **(C)** Relative quantification TopFluor-TMR-PA-derived fluorescence intensity co-localizing with lipid droplets in control (*siCtrl*) and ORP5-depleted (*siORP5*) HeLa cells. Data are represented as boxplots showing the interquartile range, the median and the mean (+) of *n* = 2815 - 2903 lipid droplets in three independent experiments. Statistical significance between data sets was assessed using Mann-Whitney U test, with ****p < 0.0001. **(D)** Representative confocal images, including full cell and zoomed regions, of wild-type (*WT*) and ORP5 knockout (*ORP5KO*) HeLa cells treated with 2 µM TopFluor-cholesterol (green) for 30min and stained with MitoTracker orange (mitochondria, red) and Lipidblue (lipid droplets, blue). Scale bar, 10 µm. **(E)** Relative quantification TopFluor-cholesterol-derived fluorescence intensity in lipid droplets in *WT* and *ORP5KO* HeLa cells. Data are represented as boxplots showing the interquartile range, the median and the mean (+) of *n* = 2815 - 2903 lipid droplets in three independent experiments. **(F)** Representative confocal images, including full cell and zoomed regions, of wild-type (*WT*) and ORP5 knockout (*ORP5KO*) HeLa cells treated with 250 µM oleic acid overnight and then treated with 2 µM TopFluor-TMR-PA (red) for 30min. Lipid droplets were stained with LipidTox (green). Scale bar, 10 µm. **(G)** Relative quantification TopFluor-TMR-PA-derived fluorescence intensity co-localizing with lipid droplets in *WT* and *ORP5KO* cells treated with 250 µM oleic acid overnight. Data are represented as boxplots showing the interquartile range, the median and the mean (+) of *n* = 417 - 826 lipid droplets in three independent experiments. Statistical significance between data sets was assessed using Mann-Whitney U test, with ****p < 0.0001. **(H)** Number of lipid droplets per cell in *WT* and *ORP5KO* HeLa cells treated with oleic acid overnight and TopFluor-TMR-PA. *ORP5KO* displays a reduced number of lipid droplets per cell. Data are represented as mean ± SEM of *n* = 20 - 21 cells. Statistical significance between *WT* and *ORP5KO* was assessed using Mann-Whitney U test, with ****p < 0.0001. **(I)** Lipid droplets volume in WT and ORP5 treated with oleic acid and TopFluor-TMR-PA. Data are represented as boxplots showing the interquartile range, the median and the mean (+) of *n* = 417 - 826 lipid droplets from three independent experiments. **(J)** Scatter plot of TopFluor-TMR-PA derived signal intensity and lipid droplets volume in *WT* and *ORP5KO* cells treated with oleic acid overnight. Data points, 417 - 826 lipid droplets, were fitted with simple linear regression with P value and R-squared indicated in the plots. **(K)** *Top:* Triacylglycerol (TAG) content in lipid extract isolated from HeLa WT and ORP5KO. *Bottom:* Phospholipids (PLs) content in lipid extract isolated from WT and ORP5KO HeLa cells. Data are represented as mean ± SEM of three independent experiments. **(L)** Experimental protocol: HeLa WT and ORP5 KO cells were transfected with either an ER marker (EROX-BFP, blue) for 24 h. Lipid droplets were then induced by 24 h oleic acid (OA) feeding and a TMR-PA pulse was performed 1 h before cell swelling with hypotonic medium (1:20 dilution). Confocal microscopy images of WT and ORP5KO cells expressing EROX-BFP after TMR-PA pulse and swelling. Lipid droplets were stained with LipidTod (LTox, green). Scale bars: 10 µm (full image), 1 µm (zoom). **(M)** Quantification of TopFluor-TMR-PA fluorescence intensity in the ER and lipid droplet. Data represents the lipid droplets to ER ratio TopFluor-TMR-PA fluorescence intensity plotted as boxplots showing the interquartile range, the median and the mean (+) of n = 32 - 33 cells in three independent experiments. Statistical significance between data sets was assessed using Mann-Whitney U test.

**Supplementary Figure 4:**
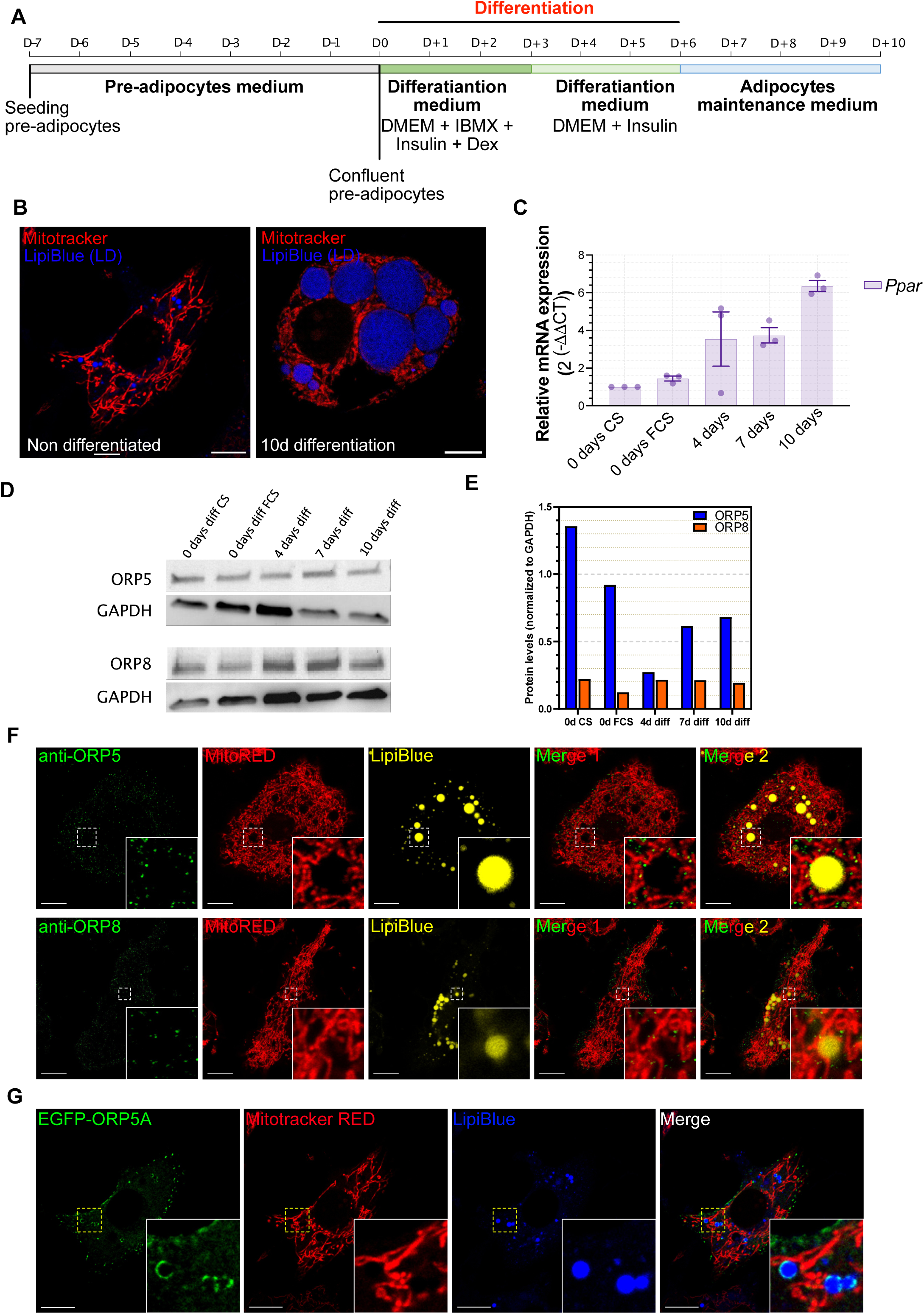
**(A)** Schematic representation of conceptual framework followed for 3T3-L1 differentiation in adipocytes. D: day, DEX: dexamethasone. **(B)** Single confocal plane representative of undifferentiated (left) and differentiated (right) 3T3-L1 cells or after 10 days of differentiation. Mitochondria and lipid droplets were labeled with Mitotracker orange (red) and LipiBlue (blue), respectively. Scale bar, 10 µm. **(C)** Expression of PPARγ by RT-qPCR. Gene expression was quantified using the 2(-ΔΔCt) method, using 26S as housekeeping gene. **(D)** Immunoblot showing the expression of ORP5 and ORP8 during 3T3-L1 cell differentiation. **(E)** Relative quantification of ORP5 and ORP8 protein expression, obtained by normalizing ORP5 expression by the housekeeping GAPDH. **(F)** Confocal image of 3T3-L1 cells after 10 days of differentiation. ORP5 and ORP8 proteins were labeled using anti-ORP5 (green) and anti-ORP8 (green) antibodies, respectively. Mitochondria and lipid droplets were labeled with Mitotracker orange (red) and LipiBlue (yellow), respectively. The insets show zoomed-in views of the circled regions **(G)** Confocal image of 3T3-L1 cells transfected with the EGFP-ORP5A (green) after 7 days of differentiation. Mitochondria and lipid droplets were labeled with Mitotracker orange (red) and LipiBlue (blue), respectively. The insets show zoomed-in views of the circled areas. Scale bar, 10 µm

**Supplementary Figure 5:**
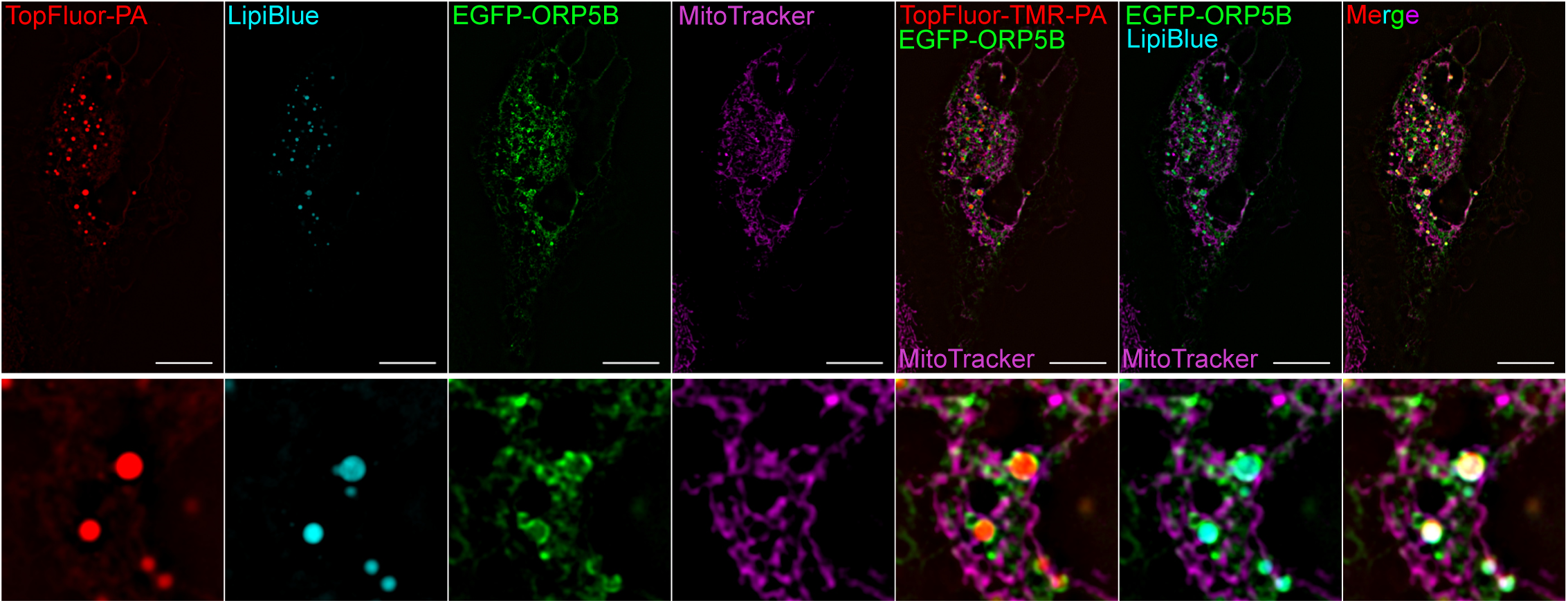
Structure eliminated microscopy (SIM) micrographs of HeLa cells expressing EGFP-ORP5B (green) or treated with TopFluor-TMR-PA (red) for 30 min, and stained with Mitotracker (mitochondria, purple) and Lipiblue (lipid droplets, cyan). Scale bar, 10 μm.

**Supplementary Figure 6:**
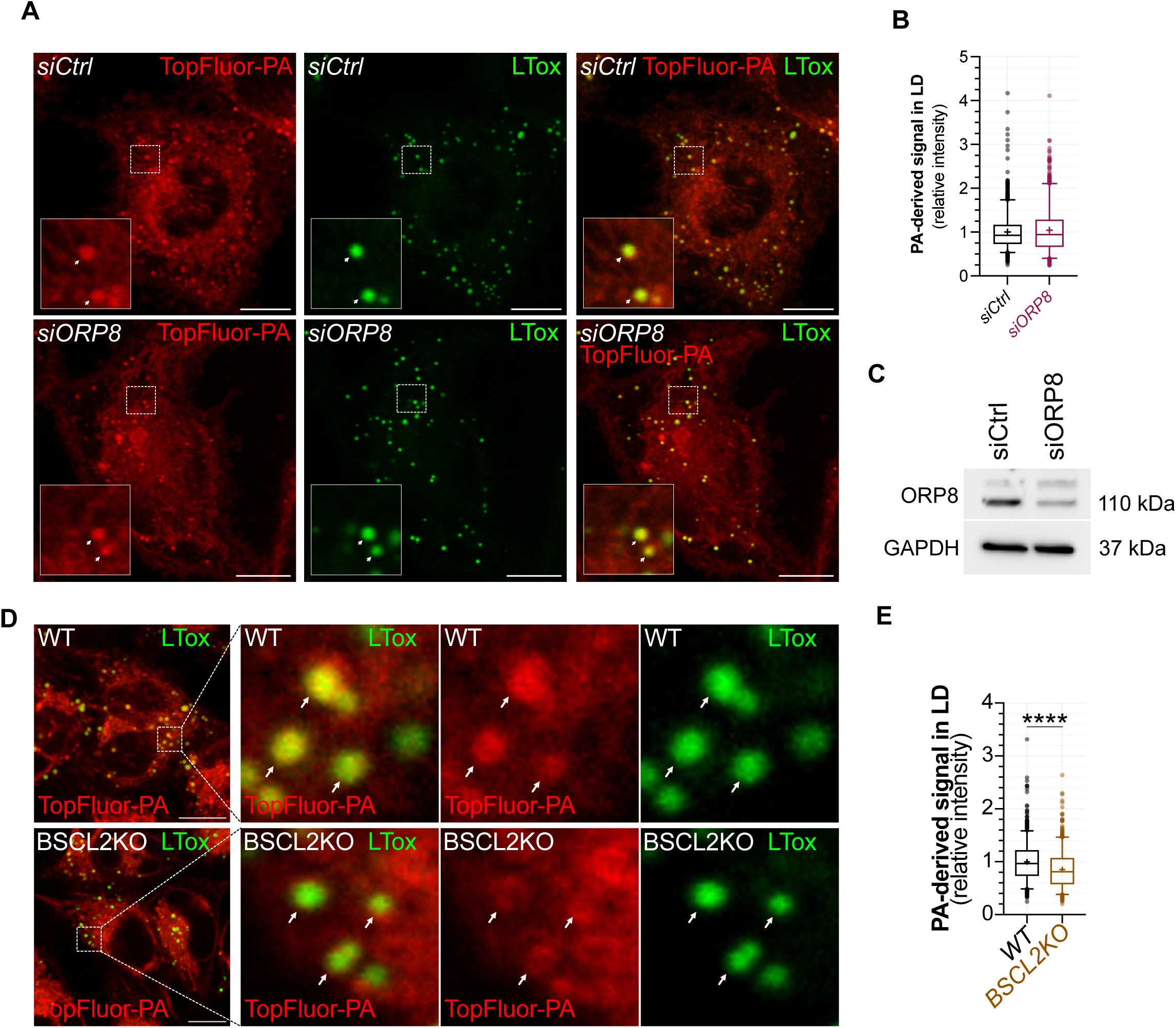
**(A)** Representative confocal images, including full cell and zoomed regions, of HeLa cells treated with 2 µM TopFluor-TMR-PA (red) for 30min 48 h after transfection with control siRNA or with a *ORP8*-specific siRNA. Lipid droplets were stained with LipidTox (green). Scale bar, 10 µm. **(B)** Relative quantification TopFluor-TMR-PA-derived fluorescence intensity co-localizing with lipid droplets in control and ORP8-depleted cells expressing. The absence of ORP8 does not affect the incorporation of TopFluor-TMR-PA in the Lipid droplets. Data are represented as boxplots showing the interquartile range, the median and the mean (+) of *n* = 946 - 1024 lipid droplets. **(C)** Representative immunoblotting confirming the efficiency of RNAi treatment to decrease ORP8 protein expression in HeLa cells. **(D)** Representative confocal images, including full cell and zoomed regions, of *WT* and *BCSL2KO* HeLa cells treated with 2 µM TopFluor-TMR-PA (red) for 30min with lipid droplets marked by LipidTox (green). Scale bar, 10 µm. **(E)** Relative quantification TopFluor-TMR-PA-derived fluorescence intensity in the lipid droplets in *WT* and *BSCL2KO* cells. Depletion of BSCL2 reduces the incorporation of TopFluor-TMR-PA in lipid droplets. Data are represented as boxplots showing the interquartile range, the median and the mean (+) of *n* = 703 - 874 lipid droplets. Statistical significance between data sets was assessed using Mann-Whitney U test, with ****p < 0.0001

**Supplementary Figure 7:**
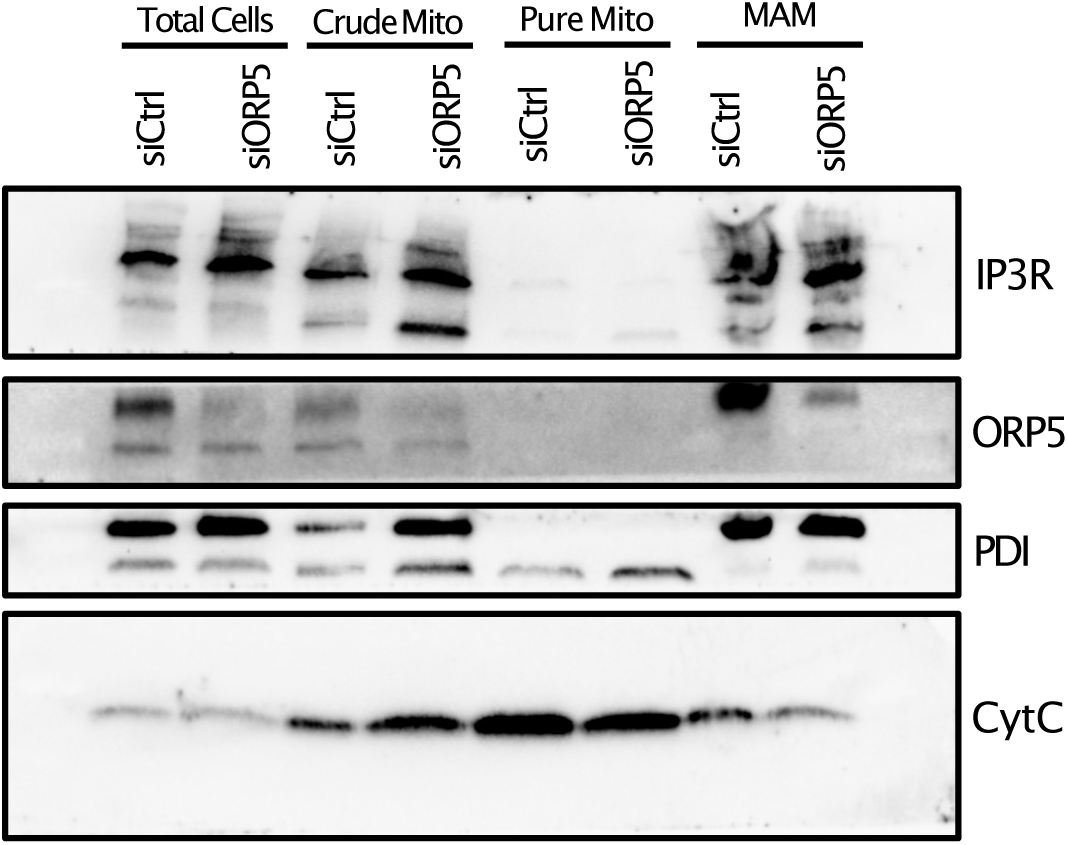
Immunoblot analysis of mitochondria (Mito) and mitochondria-associated ER membranes (MAM) proteins in subcellular fractions. Proteins analyzed include ORP5 (ER and MAM), IP3R-3 (MAM protein), PDI (ER) and cytochrome c (mitochondrial protein).

**Supplementary Figure 8:**
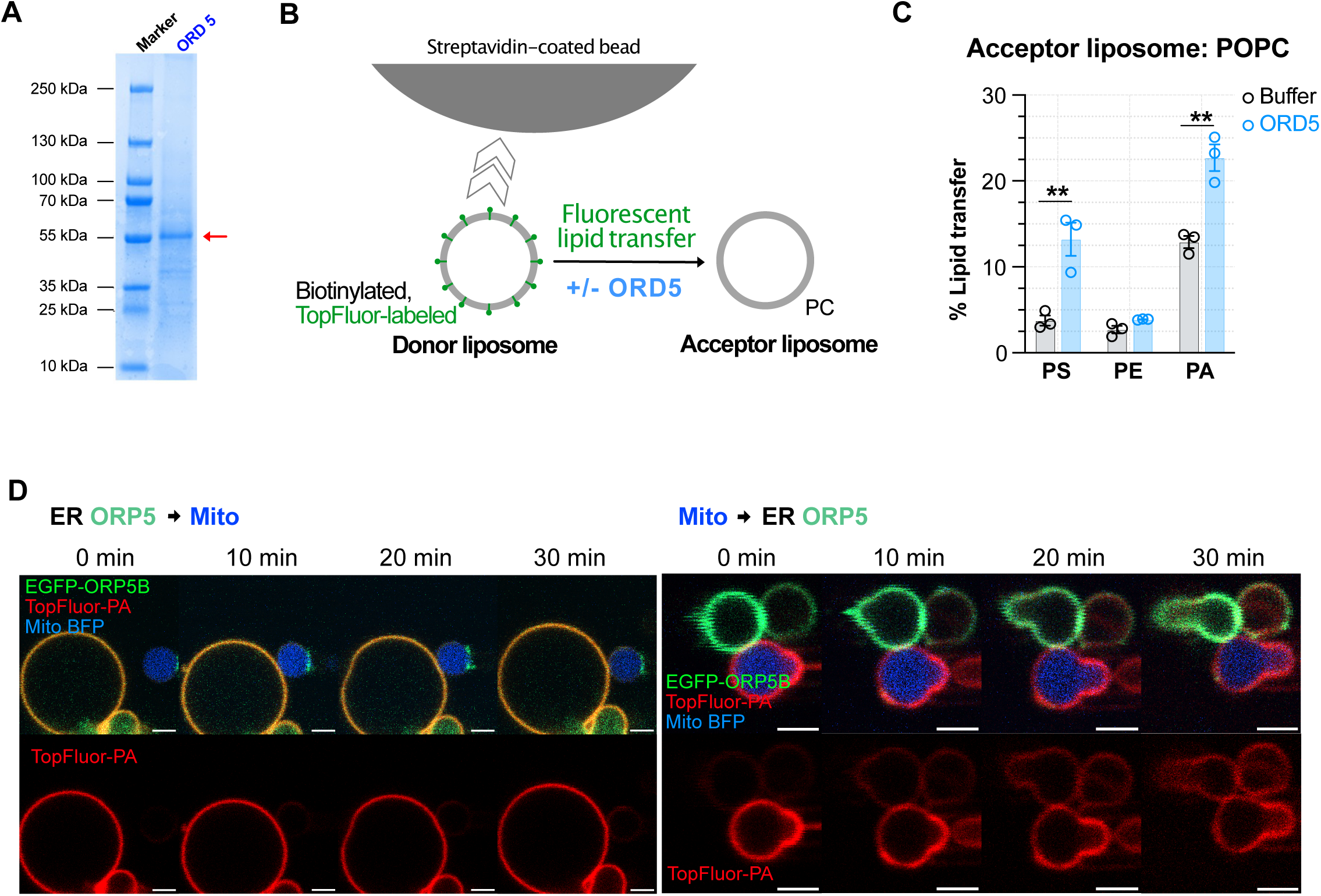
**(A)** Coomassie stained SDS-PAGE of the purified recombinant ORD domain of ORP5 protein. **(B)** Schematic representation of the conceptual framework of *in vitro* lipid transfer assay between liposomes mediated by the ORD domain of ORP5 (ORD 5). Donor Liposomes contain POPC, Biotinylated and TopFluor phospholipids. Acceptor Liposomes are composed of POPC. **(C)** Percentage of transfer of TopFluor phospholipid phosphatidylserine (PS), PE and phosphatidic acid (PA), mediated by ORD5 between the donor and the acceptor liposomes. Data are presented as % of transferred lipid ± SEM and are the mean of three independent experiments. Statistical analysis was performed using unpaired student’s *t*-test, **p < 0.01. **(D)** Donor GOVs (either giant ER vesicles (GERV) or giant Mitochondria vesicles (GMV)) fed with TopFluor-TMR-PA (red) combined with acceptor GOVs and brought into contact via micropipettes. PA transfer between GOVs upon artificially established contact, comparing representative confocal images at 0 min, 10 min, 20 min and after 30 min for the donor→acceptor pairs: ER ORP5 → Mito (left) and Mito → ER ORP5 (right), with TopFluor-TMR-PA transferring from donor to acceptor GOVs. Scale bar, 2 µm.

**Supplementary Figure 9:**
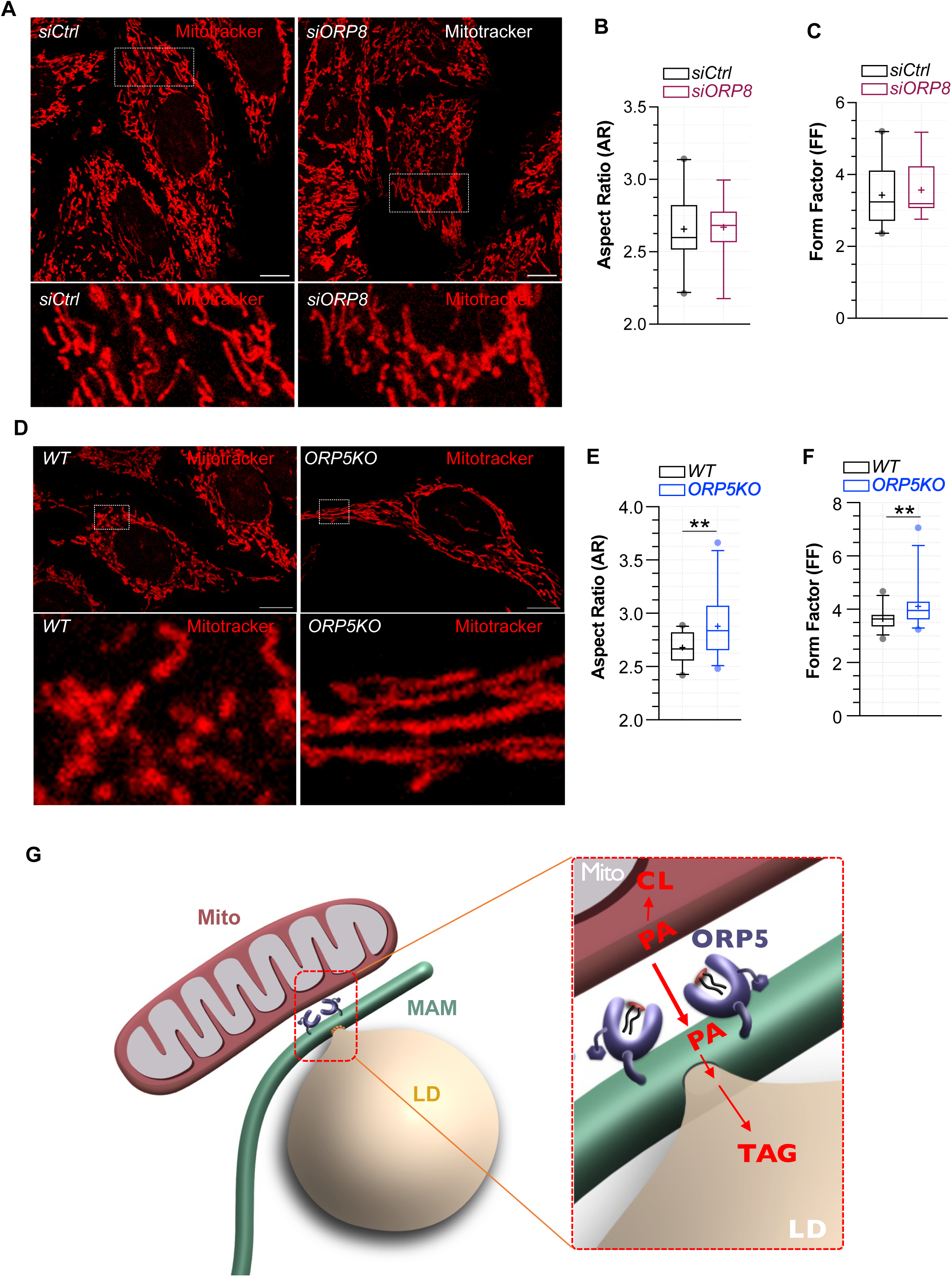
**(A)** Confocal images showing mitochondria morphology in control (*siCtrl*) and RNAi ORP8-depleted (*siORP8*) HeLa cells stained with mitotracker orange (red) to mark mitochondria. **(B-C)** Quantification of the mitochondrial shape descriptors AR and FF. Depletion of ORP8 does not affect mitochondrial morphology in HeLa cells. Data are represented as boxplots showing the interquartile range, the median and the mean (+) of *n* = 16 - 20 cells in three independent experiments. **(D)** Representative confocal images, including full cell and zoomed regions, showing mitochondria morphology in *WT* and *ORP5KO* cells **(E-F)** Quantification of the mitochondrial shape descriptors aspect ratio (AR) and form factor (FF). *ORP5KO* cells display elongated and more branched mitochondria as compared with *WT* cells Data are represented as boxplots showing the interquartile range, the median and the mean (+) of *n* = 27 - 29 cells in three independent experiments. Statistical analysis was performed using Mann-Whitney U test, **p < 0.01. **(G)** Cartoon illustrating our model: at the MAM–LD interface, ORP5 redirects PA away from excessive cardiolipin synthesis toward TAG formation, thereby coupling lipid storage with mitochondrial lipid homeostasis. This provides a novel safeguard mechanism that protects mitochondria from cardiolipin overload.

